# Six million years of vole dental evolution driven by tooth development

**DOI:** 10.1101/2025.02.24.639961

**Authors:** Fabien Lafuma, Élodie Renvoisé, Julien Clavel, Ian J. Corfe, Gilles Escarguel

## Abstract

Morphological change occurs over macroevolutionary timescales under the action of natural selection and genetic drift combined with developmental processes shaping organogenesis^1–3^. Although determining their relative weight is made difficult by discrepancies between palaeontological and neontological data^4^, mammalian tooth morphology may bridge the gap between fossil record and laboratory observations. Fossils indicate that placentals and marsupials diversified after evolving molars bearing more cusps^5–7^, which emerge through the iterative signalling of enamel knots^8,9^. However, this theoretical evo-devo model of mammalian tooth evolution has not been explicitly assessed on empirical data. Here, by combining the fossil record with laboratory experiments, we identify a shared developmental basis for the convergent, ratcheted evolution of increasingly complex molars in arvicoline rodents (voles, lemmings, muskrats). Longer, narrower molars lead to more cusps throughout development and deep time, suggesting a developmental driver for tooth evolution. Both the arvicoline fossil record and vole tooth development show slower transitions toward the highest cusp counts. This newly identified pattern suggests that the developmental processes fuelling the evolution of increasingly complex molars may also limit the potential for further complexity increases. Integrating palaeontological and developmental data shows that long-term evolutionary trends can be accurately explained by the simple tinkering of developmental pathways.

---

Descent with modification under natural selection and drift has led to the evolution of the vast morphological diversity seen throughout the history of life^1,3^. The production of variation and its sorting generation after generation are thus central evolutionary processes^4,10^. The resulting patterns of morphological change in deep time have been historically documented as gradual transitions explained by the differential success of variants under environmental pressures^1,11,12^. However, comparative developmental biology later revealed the influence of ontogenetic processes in directing morphological variation in deep time^2,4,13,14^. In contrast to previous theories, these findings underscored the capacity of mechanisms that generate variation to drive evolutionary trajectories, including large and abrupt evolutionary saltation previously dismissed as artefacts of the fossil record^4,15,16^. As such, evolutionary developmental biology offers developmental explanations for phenotypic evolution in deep time based on regulatory changes involving conserved developmental genes and pathways^17,18^, developmental constraints^19,20^, and the modulation of developmental sequences^13,21^. Because these developmental properties non-randomly shape the variation available for selection^2^, they are expected to have implications for the evolution of clades in deep time^10^. Determining the importance of this push of development relative to the pull of the environment in driving macroevolutionary trajectories thus requires combining approaches of evolution at multiple spatiotemporal scales^10,22,23^, which has been hindered by the difficult integration of palaeontological and neontological data^10,24^. In this context, studying mammalian teeth offers an opportunity to bridge this methodological gap. In the Cenozoic, placental and marsupial mammals diversified as multiple lineages independently evolved quadritubercular molars via the addition of a neomorphic cusp—the hypocone^5,6^. During tooth development, cusps are preceded by precursor enamel knots acting as signalling centres responsible for crown patterning^25,26^. The final number and placement of enamel knots—and, therefore, cusps— depends on a reaction–diffusion system of activator and inhibitor molecules modulated by a regulatory network of dental genes. It has therefore been possible to establish an evo-devo model of mammalian tooth morphology linking morphological innovations in deep time with their developmental origins^8,9,27–29^. Under the expectations of evo-devo, the regulation of existing gene networks and modulation of developmental sequences should provide preferential avenues for limited genotypic change to produce substantial phenotypic change that shapes macroevolutionary trajectories. However, currently available in vitro and in silico data point to the difficulty of increasing dental complexity during development^28,30^. Because mammalian tooth evo-devo research has focused mainly on the domestic mouse (Mus musculus), studying other taxa could reveal novel insights into the developmental origin of new tooth morphologies in deep time. Here, we demonstrate that the molar teeth of arvicoline rodents (voles, lemmings, muskrats) are a promising system for investigating the contributions of developmental push and environmental pull on phenotypic evolution. On one hand, Arvicolinae have an excellent Plio-Pleistocene fossil record documenting dental trends at high resolution over most of the Northern Hemisphere during the Quaternary glaciation cycles^31^. On the other hand, voles share a relatively recent common ancestor with mice^32^ and can be bred in laboratory conditions, allowing comparative studies to test developmentally-informed hypotheses regarding their dental evolution. Since the origin of Arvicolinae ca. 6 million years ago (Ma), their first lower molar (m1) evolved an increasingly complex occlusal surface bearing more cusps as its size increased while arvicolines diversified into the third most speciose group of muroid rodents^31–33^. Hence, the arvicoline m1 provides parallels to Cenozoic mammalian tooth evolution while also having reached extremes in tooth complexity unseen in all but one other rodent and approximating morphologies seen in the largest herbivorous mammals^30,34^. In this study, we address the evolution of the arvicoline m1 using phylogenetic comparative methods and experimentally uncover a putative simple developmental control on its morphology in deep time, providing evidence for a strong, sustained developmental push on phenotypic evolution in a time of intense environmental selection pressures.

## Results

### Convergent increases in molar complexity

Because of the evidence of increasing complexity in the m1 of Arvicolinae over time^31,33,35^, we sought to test the pattern of evolution of m1 cusp number taking into account arvicoline phylogeny. We used Maximum Likelihood ancestral character state reconstruction on a large dataset and tree of fossil and extant taxa (SI Data 1, 2) and found a general trend of increasing complexity of the m1 throughout arvicoline evolution (Fig. 1). Starting from an ancestral five-cusped tooth, the direct evolution of a seven-cusped pattern without six-cusped intermediate stage was followed by multiple independent acquisitions of increasingly complex molars with up to eleven cusps, mostly via events of double cusp addition (Fig. 1a–h). In contrast, there were few instances of single cusp additions and even fewer cusp loss events (one loss for every four gains) (Extended Data Fig. 1). According to the best-supported model of character evolution (Fig. 1h, Extended Data Fig. 2), the rates of double cusp additions towards seven and nine-cusped teeth were highest, while increases towards more complex molars (*i.e.*, ten and eleven cusps) occurred at rates one order of magnitude lesser. Consequently, arvicoline evolution followed a pattern of successive waves of abundance of increasingly complex first lower molars, starting from five cusps and progressively replaced by new morphologies bearing more cusps via two-cusp stepwise increases until reaching the currently dominant nine-cusped configuration. However, each successive replacement took more time to initiate and complete than the previous (Fig. 1i). Renvoisé *et al.*^33^ proposed that increases in m1 length during the evolution of Arvicolinae could have caused the increases in crown complexity of that tooth. Indeed, recent works have shown that shape-independent tooth scaling is actively regulated throughout tooth formation^36^, suggesting that changes in molar size without equivalent scaling of cusp size could be responsible for the lack of a six-cusp stage and the further increases in molar complexity.

**Fig. 1.**
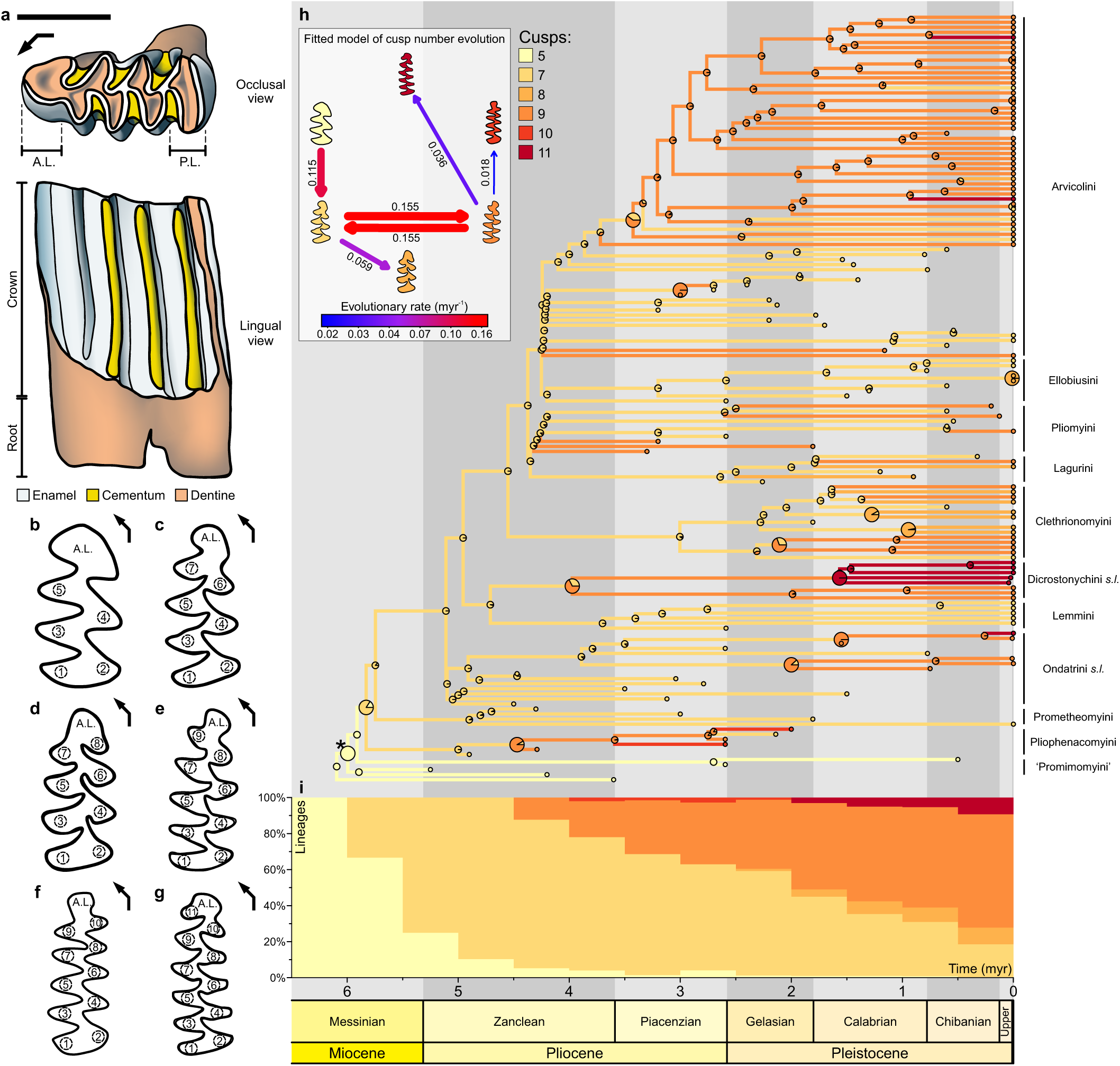
Evolution of highly complex first lower molars in Arvicolinae. **a**, Anatomy and histological composition of a right first lower molar of bank vole (*Clethrionomys glareolus*) in occlusal (top) and lingual view (bottom), after Renvoisé *et al.*^37^. Scale bar = 1 mm. Arrow: anterolingual direction. **b–g**, Schematic representation of the levels of occlusal complexity identified in living and extinct Arvicolinae, with individual cusps posterior to the anterior loop numbered from most posterior to most anterior (including two cusps forming the posterior loop; see Methods): **b** five cusps, **c** seven cusps, **d** eight cusps, **e**, nine cusps, **f**, ten cusps, **g**, eleven cusps. Not to scale. A.L.: anterior loop. P.L.: posterior loop. Arrows: anterolingual direction. **h**, Observed first lower molar cusp numbers (tips) and maximum likelihood ancestral state estimations at nodes (pie charts) across a phylogeny of 151 species of Arvicolinae (82 living and 69 extinct) based on the fitted relative rates of character transition for the best-supported model of trait evolution (inset). Larger pie charts mark the most ancient nodes with >50% relative likelihood for a given cusp number and lineage, corresponding to independent increases in cusp number. Branch colours denote character states between nodes. Ondatrini *s.l.*: Ondatrini *sensu lato* (including *Ondatra*, *Neofiber*, and related stem-taxa). Dicrostonychini *s.l.*: Dicrostonychini *sensu lato* (including *Dicrostonyx*, *Arborimus*, and *Phenacomys*). Asterisk: most recent common ancestor (MRCA) of Arvicolinae. **i**, Relative proportions of lineages bearing five-, seven-, eight-, nine-, ten- and eleven-cusped first lower molars across 500,000-year time bins over the last 6.5 million years based on our maximum likelihood ancestral reconstructions of first lower molar cusp number.

### Long, narrow teeth drove m1 complexity

To test the role of size in the increased complexity of m1, we modelled the relation of molar dimensions (length and width) and their interaction on cusp number during arvicoline evolution by fitting multiple phylogenetic and non-phylogenetic Poisson regressions. We found most relative support was concentrated by a single non-phylogenetic Conway–Maxwell–Poisson (CMP) model describing an association between cusp number and relative m1 elongation (Extended Data Fig. 3). This same model was favoured by phylogenetic and non-phylogenetic Poisson regressions with similar results (Extended Data Fig. 3a). Phylogenetic Poisson regressions were the least supported, showing no effect of phylogeny on the relationship between cusp number and molar dimensions. Similarly, phylogenetic and non-phylogenetic logistic regressions on the subsample of species bearing molars with nine and seven cusps (the most and second-most abundant m1 morphologies; SI Data 1, 3) provided the strongest relative support to cusp number correlating positively with length and negatively with width (Extended Data Fig. 4). Like for Poisson regressions, a single non-phylogenetic binomial logistic regression was largely favoured, in agreement with less supported regressions simultaneously estimating similar model parameters and weak phylogenetic signal (see Methods). Hence, while the increasing length of the m1 could control the evolution of more cuspidate molar crowns, the evolution of narrower teeth also played a critical role in the advent of more complex morphologies. Accordingly, ancestral state reconstructions of molar dimensions under a Brownian Motion process show that, throughout Arvicolinae, various trajectories that included increasing length or decreasing width relative to length led to the convergent evolution of chiefly nine-cusped morphologies, meaning some clades reached higher cusp counts without increasing molar size (Extended Data Fig. 5, SI Data 4). Increasing length was, therefore, sufficient but not mandatory to increase m1 complexity. However, we inferred an episode of marked isometric size increase of the m1 at high rates of evolution coinciding with the early diversification of Arvicolinae (Extended Data Fig. 6a–c). This might have been key for the direct transition from five to seven cusps ancestral to all further events of cusp additions without a six-cusp stage. Alternatively, substantial changes to tooth development before the radiation of crown-Arvicolinae could have driven the transition to seven cusps and the switch to a two-cusp stepwise mode of molar complexity increases. However, such a scenario appears less likely given the apparent difficulty of generating additional cusps during mammalian tooth development^28^.

### Early growth spurt of vole tooth cusps

We investigated the role of developmental processes—in particular, heterochronic shifts—in the evolution of the highly complex m1 of Arvicolinae by comparing the process of crown patterning at various stages in the developing m1 of bank voles (*Clethrionomys glareolus*) and the less cuspidate m1 of mice (*Mus musculus*) imaged using X-ray computed microtomography scanning^37^. Though both species initiate crown patterning at the same time, with their protoconid appearing at embryonic day 15 (E15), the following 24 hours show an early burst of cusp additions in the bank vole (Fig. 2a). Previous data^8^ for the East European vole (*Microtus levis*)—a species with a higher m1 cusp count (9) than the bank vole—revealed an even more pronounced burst with faster and more numerous cusp additions (Fig. 2b). This difference—and not earlier cusp initiation—allows voles to grow more cusps quicker and to reach higher final cusp counts than mice. Following this early burst of cusp additions, the rate of cusp appearance progressively decreases in voles, whereas it ramps up in mice so that, by E17, cusp growth becomes faster in mice than voles. Despite cusp additions accelerating later in mouse development, this is too slight a change and too close to the end of crown patterning to result in high cusp numbers in mice. In contrast, the heterochronic acceleration towards an early burst of cusp additions in vole molars allows for more cusps to grow despite a constantly decreasing rate of cusp appearance from that point on. These increasingly slower cusp additions mirror the evolution of the m1 of Arvicolinae, in which the first increases in cusp number (five to seven and seven to nine) occur rapidly, but the rates of transition towards the most complex morphologies (ten and eleven cusps) are substantially lower (Fig. 1h). Furthermore, the stepwise addition of multiple cusps is a staple of both the evolution and development of the m1, providing a developmental basis for the morphological evolution of that tooth that sets Arvicolinae apart from other muroids. Though heterochronic shifts are responsible for supernumerary cusps in both mice and voles^27,28,38^, the latter uniquely rely on developmental acceleration rather than the advancement of crown patterning to reach more complex morphologies, revealing distinct modes of regulation for increasing cusp number during development in voles and mice.

**Fig. 2.**
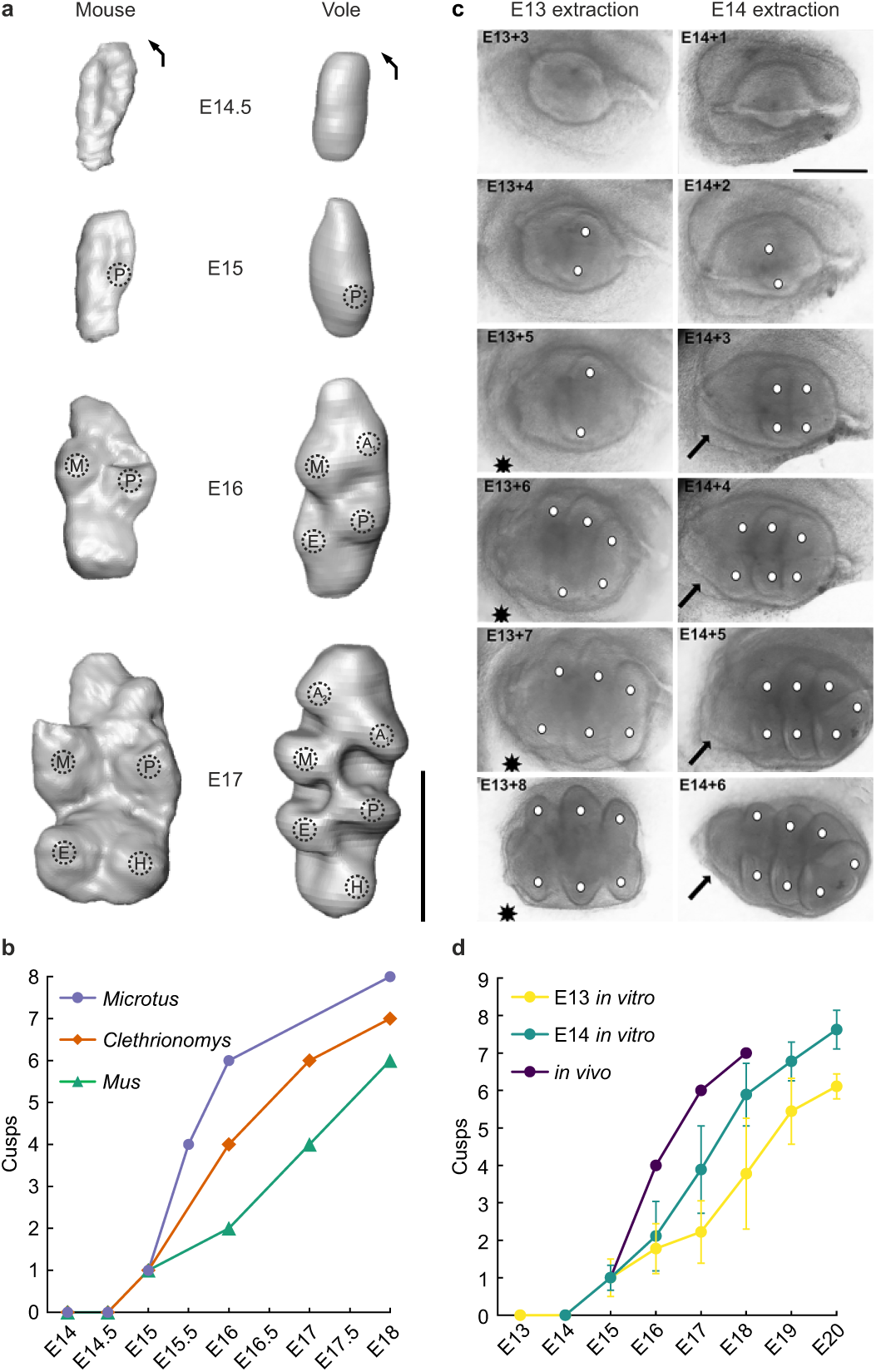
Developmental processes generating the highly complex first lower molars of Arvicolinae. **a**, Three-dimensional renderings of developing right first lower molars of mice (left) and bank voles (right) in occlusal view generated using X-ray computed microtomography scans (microCT-scans), after Renvoisé *et al.*^37^. P: protoconid. M: metaconid. E: entoconid. H: hypoconid. A_1_: first anteroconid. A_2_: second anteroconid. Scale bar = 0.5 mm. Arrows: anterolingual direction. **b**, Cusp counts in m1 germs of mice (*Mus musculus*), bank voles (*Clethrionomys glareolus*), and East European voles (*Microtus levis*) developing *in vivo* from embryonic day 14 (E14) to 18 (E18). Data for *Microtus levis* and E18 mouse m1 were sourced from Jernvall *et al.*^8^ and Christensen *et al.*^36^, respectively. **c**, Developmental sequences of bank vole left first lower molar germs extracted at embryonic day 13 (E13, left) or embryonic day 14 (E14, right) and cultured *in vitro*, photographed in occlusal view. The day of culture is noted in each frame, from one (+1) up to eight days (+8) after extraction. White discs indicate cusps. Note that the anterior part of the molar fails to develop in germs extracted at E13 (star) but is present in germs extracted at E14 (arrow). Scale bar = 0.5 mm. **d**, Cusp counts in first lower molar germs of bank voles developing *in vivo* (see Fig. 2b) and *in vitro* after extraction at E13 (*n* = 72) or E14 (*n* = 61). Vertical bars = ±1 standard deviation.

### Tooth length controls molar cusp count

Arvicolinae showed a strong correlation between molar dimensions and cusp number during their evolution. However, experimentally inducing supernumerary cusps in mice leads to more tightly packed cusps without substantially increasing molar size^28^. To characterise the developmental basis of the observed evolutionary increases in complexity, we extracted the developing m1 of bank voles one or two days before the appearance of the protoconid (*i.e.*, at E14 and E13, respectively) and cultured them *in vitro* for the remainder of the crown patterning process. An earlier extraction resulted in shorter E13 germs with fewer cusps due to these teeth failing to elongate anteriorly and develop a complete anteroconid complex and anterior loop (Fig. 2c, Extended Data Fig. 7, SI Data 5). Nevertheless, the *in vivo* dynamic of cusp additions was preserved in both experimental conditions (Fig. 2d). The time of appearance of the protoconid was not affected by the extraction protocol, which chiefly delayed the initiation of the burst of cusp additions. This delay was longer, with fewer subsequent cusp additions in germs extracted at E13, while multiple E14 germs reached the normal final cusp count for the bank vole m1 (*i.e.*, eight cusps). Therefore, a key event (or suite of events) taking place between E13 and E14 determines the normal progression of the early spurt of cusp addition characteristic of voles. Fitting and comparing various CMP regression models on these data revealed that, while cusp number positively correlates with length and width in our experiments, increasing width strongly diminishes the effect of length (Extended Data Table 1, Extended Data Fig. 8). Consequently, only E14 germs underwent sufficient growth in length to experience sustained cusp additions; despite being cultured for one more day, all E13 germs failed to recover the normal m1 final cusp count. Notably, we observed the simultaneous appearance of two adjacent cusps (one lingually, one buccally) in both populations of molar germs (Fig. 2c), and all germs experienced at least one step of multiple (2-4) cusp additions during their development (SI Data 5). Though double cusp additions did not occur throughout molar development nor in all germs, they suggest an equivalent of macroevolutionary two-cusp stepwise transitions at the developmental level. Remarkably, our culture experiments demonstrate a relationship between molar dimensions and cusp number paralleling that observed in the evolution of Arvicolinae, despite molars developing *in vitro* growing into markedly different proportions from the *in vivo* conditions due to the absence of mechanical support from the jawbone^37^. Meanwhile, the different effects of width in each dataset (*i.e.*, a negative effect on cusp number during evolution and a positive impact during development) can be attributed to the developmental regression model describing the growth of the molar from a small-sized germ into a larger-sized tooth, while macroevolutionary data included only fully developed adult teeth. Nevertheless, the negative interaction of length and width was even stronger during development than throughout evolution, emphasising the importance of limiting the lateral expansion of the molar to increase cusp number at both scales. Furthermore, as earlier extraction makes the pattern of cusp additions closer to our observations of the mouse molar (*i.e.*, a longer initial period of slow cusp addition followed by an acceleration), the heterochronic shift toward accelerated cusp development characteristic of voles was likely caused by a derived developmental condition manifesting between E13 and E14 that evolved only in Arvicolinae, resulting in their uniquely complex m1.

## Discussion

From these data on vole molar development, we can formulate hypotheses regarding the nature of the developmental processes that both determine the early burst of cusp additions and ensure the elongation and complete patterning of the tooth germ in molars extracted at E14 rather than E13. Previous works proposed that the molar placode merging with a vestigial premolar germ could explain the anterior elongation of the m1 of Arvicolinae^38,39^. In mice, a signalling centre (R2) transiently appears anteriorly to the m1 germ between E13 and E14^40,41^. R2 then fuses with the m1, which elongates^40^. Because the timing of protoconid initiation is conserved between mice and voles, extraction at E13 would have preceded the fusion of a putative R2-like centre with the m1 germ, possibly explaining the shorter molars generated in that experimental condition. However, there is no evidence that such a structure exists in voles. Most importantly and contrary to voles, in mice the fusion of R2 results in a narrow anterior extension of the m1 and a decrease in occlusal complexity due to lesser activation signalling^41^, implying substantial changes in a putative vole equivalent of that structure. Alternatively, the mechanical constraints of the surrounding jawbone—which are necessary for the elongated shape of the m1 to develop^37^—could have been co-opted to generate simultaneously longer, narrower molars and more cusps by increasing the distance between enamel knots. Indeed, mechanical action is sufficient to generate the arvicoline cusp offset in voles and mice *in vitro*, effectively outsourcing aspects of crown patterning to the jawbone^37^ and possibly lowering the threshold for cusp additions, similar to how simple changes to growth biases enabled the evolution of hypocones in therian upper molars^7^. Despite the mechanical constraints of the jawbone resulting in molar elongation and thus higher cusp counts, their absence in our *in vitro* cultures did not prevent germs from showing a burst of cusp additions nor reaching normal or nearly normal final cusp counts. Hence, the dynamic of cusp development is robust to extraction from the jawbone and determined as early as E13, two days before the protoconid appears. Experimental data and *in silico* models of tooth development point to the role of increased activation signalling in increased crown complexity^7,28,30,38^. In mice, the overexpression of developmental pathways responsible for activation signalling (*e.g.*, EDA, activin A) or molecules inhibiting inhibitory pathways (*e.g.*, cyclopamine) greatly increases the number of cusps by advancing their development^28,38^. Similarly, explorations of simulated molar developmental landscapes showed that high activation is required to generate the most complex morphologies, with increased activation enabling the fastest routes along the few paths leading to them^7,30^. While these results point to intense activation signalling critically affecting vole molar development, their unique process of crown patterning likely relies on additional factors. Specifically, beyond a certain threshold, simulations show that higher activation leads to unviable tooth morphologies rather than more cusp additions^30^, implying that substantial changes to tooth developmental pathways are needed to generate the most complex molar morphologies.

In addition to our findings of a unique developmental regulation of molar complexity in Arvicolinae, comparative studies point to their m1 being among the most complex of all rodents, even approaching morphologies seen only in the largest herbivorous mammals^30,34^. Based on empirical data and the exploration of simulated developmental landscapes, Burroughs^30^ proposed that the maximum cusp number for the rodent m1 is phylogenetically constrained at nine cusps. Changes to activator and inhibitor levels in simulations did not produce viable morphologies bearing more cusps, and only the capybara (*Hydrochoerus hydrochaeris*) was found to break that limit at eleven cusps. Though similar data on the mammalian m1 in general is lacking, a study of upper second molars in placentals found that elongated teeth with multiple lateral pairs of cusps (similar to arvicoline molars) evolved only in a limited number of placental lineages^7^. Despite simulations showing this molar morphology accumulates the most cusps—under the prerequisite of high activation signalling—only a few of these taxa evolved more than nine cusps. In contrast, nine-cusped first lower molars are the most widespread among Arvicolinae, having evolved at least 19 times, and more complex teeth bearing ten or eleven cusps evolved at least two and four times independently (respectively) over the last three million years (myr). Arvicolinae thus appear to have broken away from constraints on rodent and mammalian maximum molar complexity. This departure from the dominant macroevolutionary pattern of mammalian molar variation can be explained by the heterochronic early burst of cusp development of voles. Though uniquely distinguishing voles from other muroids, it nonetheless most likely relied on a shared process of increased activation signalling that drives dental complexity across mammalian developmental landscapes, especially as activator-related parameters may have a stronger influence on increases in dental complexity than changes in inhibition signalling^7^. Importantly, the convergent evolution of molars with nine cusps or more shows that Arvicolinae repeatedly freed themselves of the constraints on molar complexity. As modulations of the same developmental process (*i.e.*, acceleration) control molar cusp counts in the two species of voles we investigated *in vivo*, representing two separate clades (Clethrionomyini and Arvicolini), we suggest it is shared by at least crown-Arvicolinae and that its potential for unlocking extreme amounts of molar complexity was frequently tapped throughout the group’s history.

In line with the uniqueness of arvicoline m1 complexity, Arvicolinae also do not follow the inhibitory cascade (IC) model, which governs molar proportions for most extinct and living mammals^33,42^. This is again due to their elongated m1, which over-dominates the other cheek teeth in relative size^33^. Because m1 length and complexity positively correlate, the same developmental processes controlling the increasing molar complexity likely contributed to the unique regulation of molar sizes in Arvicolinae. Accordingly, we found that a longer and narrower m1 evolved convergently (Extended Data Fig. 5), implying that there were multiple departures from the IC or IC-like conditions that explain the gaps in molar proportions reported across the group^33^. Hence, we suggest the same derived developmental process led to Arvicolinae simultaneously breaking away from the dominant macroevolutionary patterns for both mammalian molar complexity and size.

Despite the particular changes that pushed dental complexity to extremes in Arvicolinae, their m1 remains illustrative of the shared trends in mammalian dental evolution. Over the past 66 myr, therian mammals (*i.e.*, placentals and marsupials) have repeatedly evolved more complex molar morphologies via the addition of new cusps, most notably the hypocone. This drove the radiation of herbivorous clades with cheek teeth suited for feeding in the open grasslands that spread as the Cenozoic climate cooled^5,6,43,44^. Meanwhile, evolutionary losses of cusps were scarce^7,45–48^, unlike in other tetrapods such as squamates, whose dental evolution was labile with numerous instances of both increases and decreases in occlusal complexity^49^. Furthermore, non-reversing ratchet-like trends in mammalian tooth evolution have been observed in relation to dietary specialisation^44,50,51^. Under successive intervals of rapid evolution driven by intense selective pressures alternating with stasis as these pressures diminish, morphologies adapted for extreme conditions tend to persist rather than revert. This dynamic fosters a stepwise, irreversible progression of specialised traits, which often outlast the environmental pressures that originally favoured them, as seen, *e.g.*, with the highly complex molars of proboscideans^44,52^. The evolution of m1 occlusal morphology in Arvicolinae offers parallels to all these staples of mammalian dental evolution across 6 myr. The repeated independent evolution of neomorphic cusps drove the m1 along a ratcheted trend of increasing complexity. This trend took place while Plio-Pleistocene environments cooled and dried^53,54^ and orbital, atmospheric, and tectonic forcing triggered the cycle of Quaternary glaciations^55–58^, and resulted in the diversification of Arvicolinae in Northern Hemisphere steppes^31,59^. Therefore, despite a limited temporal range, the evolution of the m1 of over 150 species of Arvicolinae provides direct insights into evolutionary trends affecting the entire history of mammalian cheek teeth, illustrates their pervasiveness over widely different scales, and offers both the palaeontological and developmental tools to model their evolution.

The numerous parallels with the evolution of therian molars suggest that the convergent and unidirectional trend toward increased m1 complexity throughout the radiation of Arvicolinae results from selective pressures towards an adaptive peak. Importantly, both *in vivo* and *in vitro* vole molar development shows that the early burst of cusp additions is a robust and conserved process whose modulation is sufficient to produce different degrees of crown complexity. Altogether, these observations indicate that a unique developmental process can explain the macroevolutionary pattern of m1 evolution in Arvicolinae. Thus, simple tweaking of early molar development (likely relying on activation signalling^7,30^) may have enhanced the evolvability of the m1, facilitating transitions towards more cuspidate teeth. Other aspects of molar development and ontogeny could have contributed to this trend. The early evolution of a flat crown surface in Arvicolinae likely relaxed constraints on occlusion and enabled the evolution of extreme cusp counts^44,60^. Similarly, we confirm the evolution of taller teeth— hypsodonty and hypselodonty—followed the initiation of complexity increases in the arvicoline m1^35,61^. This most likely created a positive feedback loop where a more complex occlusal surface allowed for a more abrasive diet but required a taller crown to maintain occlusal morphology despite increased wear^62^. Developmental processes controlling molar morphology were thus key for the adaptive response driving the increasing m1 complexity, demonstrating that developmental processes critically shape adaptive landscapes and determine pathways of traversal toward adaptive peaks^7,63,64^.

Additionally, there is evidence that developmental processes determine the rate of morphological evolution. Indeed, non-linear developmental reaction norms generate complex genotype–phenotype maps, meaning that developmental processes control the variation available to natural selection as the result of genotypic changes^2,7,63,65^. This is because phenotypic adaptation depends on the complexity of the underlying genotype–phenotype map^66,67^ which determines how easily fitness landscapes can be traversed by phenotypes ranging from protein structure^68^ to simulated tooth morphologies^66^. *In vivo* and *in silico* evidence shows that changing the expression of tooth developmental pathways (especially activation) results in non-linear reaction norms of tooth morphology heterogeneously affecting the crown^28,29^. Thus, different morphologies of developing teeth accumulate occlusal complexity at different rates, leading to the non-independence of cusp number and pattern during development^7^. Such developmental thresholds or non-linear reaction norms could explain the stepwise evolution of crown complexity in the arvicoline m1, which, although in contrast with previous works focusing on more gradual patterns of morphological change^31,38^, can still be summarised as a continuous gradual pattern (Fig. 1i). Knowledge of developmental processes is thus necessary for interpreting macroevolutionary trends of morphological evolution, especially to increase the robustness of using the arvicoline m1 as a biostratigraphic and palaeoclimatic proxy^69^.

The developmental control on molar morphological evolution might have also been strong enough to be the main driver of the increasing complexity of the m1, with a limited added selective value between successive occlusal morphologies and over-adaptation relative to environmental conditions^44,52^. This could be why simpler morphs were not completely replaced by more complex ones, perhaps because the latter increase the potential dietary range without affecting the animals’ preference for softer foods when available^70,71^. Indeed, developmentally-informed models indicate that—at least in some mammalian groups—the evolution of molars may be nearly neutral, with the only constraint on morphology being to conform with developmental processes^72^. Under this ‘fly in a tube’ model, the properties of mammalian tooth development allow for tooth morphology to evolve along a corridor of least resistance, mostly free of selective pressures^72,73^. These properties may be responsible for the non-reversing, ratcheted evolutionary trends of increasing complexity in mammalian molars^44^ and the existence of morphological complexity traps unknown in vertebrate groups with more labile tooth development^49,74^. Despite this sustained, rapid trend of increasing complexity, the rate of evolution toward the most complex arvicoline molar morphologies (*i.e.*, ten and eleven cusps) was comparatively lower. This is striking given the abundance early in the clade’s history of taxa with a nine-cusped m1, which were well-positioned to undergo this transition. This threshold is likely a direct consequence of the decrease in the rate of cusp appearance late during molar development we observed across vole species. Accordingly, models and experiments show that the most complex molar morphologies have fewer developmental and evolutionary pathways leading to them than simpler ones^7,28^. Based on the example of Arvicolinae, we thus propose that a zone of increasing resistance exists at the high-complexity end of the corridor of mammalian tooth evolution^72^, where exponentially complex changes to development are required to produce only limited increases in morphological complexity. The same developmental processes that fuelled molar evolution in mammals may also limit its morphology and adaptive potential.

The production and sorting of variation are essential processes spanning all scales from micro- to macroevolution^4,10^. However, studies of evolution in deep time more often emphasise the differential success of variants, without examining the mechanisms responsible for generating the variation subject to selection^10,16^. Consequently, discerning environmental pull and developmental push on evolutionary trajectories remains challenging. Here, we find a strong driver of morphological evolution in deep time with a simple developmental basis. Our results confirm the critical control of tooth development on mammalian molar evolution. The complex morphology of vole molars appears to result from simpler regulatory changes than expected based on mouse molar development, though some species may have reached the phenotypic limits of this developmental tinkering. The resulting unidirectional evolutionary trend shows a strong push of tooth development over the whole history of Arvicolinae. Meanwhile, ratcheted phenotypic change suggests periods of intense selection on tooth complexity alternating with stasis when environmental pressures lessened or reversed with glacial–interglacial cycles. Disentangling the joint action of developmental processes and environmental pressures on phenotypic evolution requires integrative approaches connecting the fossil record, neontological data, and laboratory experiments. Only through this complete scope of evo-devo can we simultaneously address the variation- and selection-based forces driving evolution.

## Methods

### Phylogeny

We assembled an informal super-tree^75^ including 151 species of Arvicolinae (82 living and 69 extinct, spanning 11 tribes over the entire recorded history of the clade) and one extinct cricetid (*Baranomys longidens*) as the outgroup taxon (SI Data 2). We followed the topology and node calibrations of the muroid molecular phylogeny of Steppan & Schenk^32^ for most extant species, completed with information from several other molecular phylogenies^76–78^ for the species not sampled by Steppan & Schenk. For extinct species, we followed the relationships proposed in previous works^31,79–81^, including five anagenetic lineages (*Ogmodontomys sawrockensis-poaphagus*, *Ondatra idahoensis-zibethicus/annectens*, *Dinaromys dalmatinus-bogdanovi*, *Ungaromys dehmi-nanus*, and *Mimomys occitanus-ostramosensis*)^31^. We checked the proposed relationships for congruence with the first appearance datum (FAD) and last appearance datum (LAD) available in the Paleobiology Database (accessed on 2018-04-23; https://www.paleobiodb.org) and cross-referenced with the original publication referred in the database entry and any other related works providing additional dating information regarding the entry, particularly for publications issued prior to the redefinition of the Pliocene– Pleistocene boundary^82^. In the absence of time-calibrated phylogenies of Arvicolinae including fossil taxa, we used the cross-referenced FAD–LAD data from the Paleobiology Database for the node and tip calibrations of extinct species. For extant species without fossil record (*i.e.*, no FAD) nor estimated time of divergence based on molecular data, we arbitrarily fixed the divergence from their sister species at the beginning of the Holocene (0.0117 Ma). In the case of phylogenetic relationships implying divergence times more ancient than the recorded FAD (*i.e.*, ghost lineages) of any of the concerned taxa, we placed nodes at regular time intervals in between known node calibrations while preserving the proposed relationships. In the case of anagenetic lineages, we represented chronospecies as dichotomies with a null-length branch marking the FAD of each chronospecies. The tree includes an unresolved node reflecting phylogenetic uncertainty. For taxonomic reference, we referred to the October 28^th^, 2020 version of the ITIS database (accessed on 2020-10-29; https://www.itis.gov) and the Paleobiology Database (accessed on 2018-04-23; https://www.paleobiodb.org) for extant and fossil species (respectively).

### Morphological dataset

We assembled a dataset of cusp counts for the first lower molar (noted m1) of 151 species (82 living and 69 extinct) of Arvicolinae and the extinct cricetid *Baranomys longidens* based on existing data from the literature (SI Data 1). Additionally, we used the pre-existing literature to gather or produce linear measurements of the first lower molar (length and width) for 127 of those species (67 living and 60 extinct) using ImageJ 1.53c^83^.

To determine the number of cusps on a first lower molar, we adapted the complexity ranking of Markova^84^, itself derived from the historical nomenclature for the occlusal surface of molars in Arvicolinae of Hinton and Hibbard^85,86^ (Fig. 1a–g). In agreement with these schemes, we counted each enamel prism (‘triangles’ in the historical nomenclature)—corresponding to the entoconid, protoconid, metaconid or anteroconids—as one cusp. Only complete prisms as defined by Markova^84^ (*i.e.*, where the enamel wall forms a salient angle separated from the anterior loop by a re-entrant angle) were tallied in the cusp counts while emerging prisms (*i.e.*, where only a salient angle is visible, without separation^84^) were not counted. To maximise compatibility with developmental data and particularly cusp development in mouse first lower molars, we counted the posterior loop as two separate cusps (the hypoconid, buccally, and hypoconulid, lingually) in our cusp counts (Fig. 1b–g). In the case of documented polymorphism, we limited the sampling of cusp count and measurement data to the most common m1 morphology. When linear measurement data was available for multiple individuals of a given species with the same number of cusps, we included the averaged linear measurement values for that species in the dataset (see SI Data 1 for the sample sizes of each species).

### Ancestral character state reconstructions

We used Maximum Likelihood (ML) ancestral character state reconstruction under a time-reversible continuous discrete Markov process—the extended M*k* model for discrete character evolution^87–89^—implemented in phytools 1.9-16^90^ to estimate the evolution of the number of cusps of the first lower molar for 15 nested character transition models (Extended Data Fig. 2a). We compared the relative support for each fit using the corrected Akaike Information Criterion (AICc)^91^ (Extended Data Fig. 2b) and used stochastic character mapping (1,000 simulations) to retrieve marginal ancestral character states at nodes based on the fitted character transition matrix of the transition model with highest relative support.

To reconstruct the evolution of first lower molar dimensions, we generated a second phylogeny (SI Data 4) corresponding to the original super-tree pruned to the 127 species with known m1 dimensions in our dataset (SI Data 1). We used ML ancestral state reconstructions for continuous traits under a Brownian Motion (BM) process applied to log_10_-transformed length, log_10_-transformed width, and the length/width ratio, computing ML estimates of ancestral node values via rerooting of the tree and interpolating node states along branches as implemented in the function fitMk of phytools 1.9-16^90^. For the comparison of changes in cusp number and variations in molar dimensions (Extended Data Fig. 5), we reconstructed the ancestral states for m1 cusp number at the nodes of the phylogeny of 127 species with known molar dimensions (SI Data 4) under the best-supported model identified for the 151-species phylogeny (Extended Data Fig. 2). Similarly to the discrete ancestral character state reconstruction for the full tree, we retrieved marginal ancestral states using stochastic mapping (1,000 simulations) and identified the branches with increasing or decreasing m1 cusp number (Extended Data Fig. 9).

Ancestral reconstructions on this pruned tree presented no differences with results for the full phylogeny beyond those expected due to the absence of certain taxa.

### Rates of phenotypic evolution

We used the variable rates model of BayesTraits 3.0.2^92,93^ to estimate the rate of evolution of the dimensions (log_10_-transformed length, log_10_-transformed width, and length/width ratio) of the first lower molar. This method detects shifts in the rates of evolution of continuous traits— modelled by a Brownian motion (BM) process—across the branches of a phylogenetic tree with a reversible-jump Markov Chain Monte Carlo (rjMCMC) algorithm. Through this approach, the location of rate shifts (*i.e.*, the product of a set of rate scalars with a homogeneous background rate) is estimated using two mechanisms altering either individual branches at a time or entire clades. For each run, we conserved the default gamma priors for the rate scalar parameters and uniformly scaled the entire tree to reach an average edge length of 0.1 (*i.e.*, scaling factor = 0.102) as recommended by the software manual to improve rate estimations. We confirmed rate heterogeneity by comparing the fit of the variable rates model against a null single-rate BM model. We ran the computation five times independently using default rate priors, with each run comprising four independent chains and lasting 110,000,000 iterations. We excluded the initial 10,000,000 iterations as burn-in and sampled parameters every 10,000 iterations. We verified convergence visually in each chain and checked them for sufficient effective sample sizes using coda 0.19-4^94^ for R^95^. We retrieved the marginal likelihood of variable rates and null models through a stepping stone sampler^96^ (250 stones run for 10,000 iterations) and compared them with log Bayes Factors (logBF) to measure relative support^97^ (Extended Data Fig. 6d). Lastly, we plotted the estimated rates on the entire tree for each dimension using phytools 1.9-16^90^, ggtree 3.8.0^98^, and viridis 0.6.4^99^ for R^95^, with branches colour-coded based on the rate scalars averaged across posterior samples (as returned by the Variable Rates Post Processor; http://www.evolution.reading.ac.uk/VarRatesWebPP/), indicating relative deviation from the background rate of change (Extended Data Fig. 6a–c).

### Regression analyses

We evaluated the role of the length and width of the first lower molar on its number of cusps during the evolution of Arvicolinae by fitting a candidate set of Conway–Maxwell–Poisson (CMP), Poisson, and phylogenetic Poisson regression models using Maximum Likelihood (ML) estimation^100,101^ as implemented in glmulti 1.0.8^102^ with the fitting functions glmmTMB of glmmTMB 1.1.9^103^, glm of stats 4.4.2^95^, and pglmm of phyr 1.1.0^104^, respectively. For each type of regression, the candidate set of models included all possible combinations of predictors and their interaction term and a null model (*i.e.*, eight models in total; Extended Data Fig. 3a) to the subset of our species-level data with known cusp number and m1 dimensions (*n* = 127; SI Data 1). We then calculated the corrected Akaike Information Criterion (AICc)^91^ to measure the relative support of each fit (Extended Data Fig. 3a). Although the best-supported model gathered most of all statistical support (*ω*AICc = 0.66), because the AICc value for the second best-supported model was within two units of the best-supported model, we compared the two through a *χ*² test^105,106^, which revealed no statistically significant difference (*χ*² = 0.832, df = 1, *p*-value = 0.361, *V* = 0.081). We focused on the best-supported model for the remainder of the analyses due to its gathering of the majority of all statistical support (see Extended Data Table 1 for the full reporting of parameter estimates). We tested for misspecification using Pearson’s *χ*² goodness-of-fit test^105^ (*χ*² = 127.289, df = 123, *p*-value = 0.377, *V* = 0.090). We evaluated the quality of the fit by simulating scaled randomised quantile residuals^107^ (standardised to values ranging from 0 to 1) from the fitted model (1,000 simulations) and checked for dispersion by comparing them to the observed residuals with DHARMa’s non-parametric dispersion test^108^ (*ϕ* ∼ 1, *p*-value = 0.978, two-sided). The same protocol revealed severe under-dispersion for all Poisson and phylogenetic Poisson regressions, likely explaining their weak support relative to CMP regressions (Extended Data Fig. 3a). Additionally, we fitted phylogenetic Poisson regressions using the method of Generalised Estimating Equations (GEE)^109^ with the phyloglm function of phylolm 2.6.5^110^, though the function’s lack of implementation of the AIC prevented direct comparison with the other models. While this method struggled to recover significant fixed effects, equivalents of the best (cusps ∼ 1 + L + W:L) and second-best (cusps ∼ 1 + L + W) supported phylogenetic Poisson models fitted with phyr^104^ also estimated a significant negative interaction term between length and width and a strong significant negative fixed effect of width, respectively, in line with all other regressions supporting the role of relative elongation in increasing m1 cusp number (Extended Data Table 1).

To further investigate the relationship between the dimensions of the first lower molar and cusp number, we applied the same approach focused on a subset of our data limited to species with either a seven- or nine-cusped m1 (*n* = 109), representing the second-most and most common species in our dataset, respectively (SI Data 1). First, we fitted using ML multiple logistic binomial and phylogenetic logistic regressions through the functions glm of stats 4.4.2^95^, pglmm of phyr 1.1.0^104^, and phyloglm of phylolm 2.6.5^110^, the latter function allowing to simultaneously estimate model parameters and phylogenetic signal via bootstrapping (1,000 iterations)^111^. For each class of regression, we fitted the same candidate set of eight models including all possible combinations of predictors (*i.e.*, m1 length and width) and their interaction and a null model without predictors—on this subset transformed to have a binary response (“seven cusps” = 0, “nine cusps” = 1). We compared the relative support for each fit using the AICc^91^ and found the best-supported model to gather a large majority of relative support (*ω*AICc = 0.62; Extended Data Fig. 4a). We did not include the models fitted with phyloglm due to the function currently not supporting the AIC. We checked the best model for misspecification using the Hosmer–Lemeshow goodness-of-fit test^112^ (*χ*² = 8.423, df = 8, *p*-value = 0.393) as implemented in ResourceSelection 0.3-6^113^. The full statistical reporting of parameter estimates for all models is detailed in Extended Data Table 1. Because misspecification does not result in dispersion for Bernoulli processes, we did not check the models for dispersion. Phylogenetic logistic regressions including the simultaneous estimation of model parameters and phylogenetic signal fitted with phyloglm recovered similar estimates to those of other regressions (Extended Data Table 1) with a weak signal for all models (α ranging 0.18-0.34; results not shown) except for the null model without predictors (α = 5.93). We examined the link between molar dimensions and cusp number during development by fitting to the measurement and count dataset of developing first lower molars cultured *in vitro* (*n* = 133; SI Data 5) an exhaustive candidate set of 36 CMP regressions^100,101^ (including a null model and all possible combinations of length, width, extraction day, and their first-degree interaction terms as predictors; Extended Data Fig 8a) with maximum likelihood in glmulti 1.0.8^102^. We compared the relative support for each model using the AICc^91^. Though the best-supported model gathered most of all statistical support (*ω*AICc = 0.75; Extended Data Fig. 8a), we compared it to the second-best model due to their AICc values differing by less than two units: a *χ*² test^105,106^ revealed no significant difference (*χ*² = 0.044, df = 1, *p*-value = 0.833, *V* = 0.018). We thus limited the remainder of the analyses to the best-supported model (see Extended Data Fig. 8b for the complete reporting of parameter estimates). We used Pearson’s *χ*² goodness-of-fit test^105^ to check the absence of misspecification (*χ*² = 118.029, df = 126, *p*-value = 0.681, *V* = 0.084). To check for dispersion, we simulated scaled randomised quantile residuals^107^ from the fitted model (1,000 simulations) for comparison with the standardised observed residuals in the non-parametric dispersion test of DHARMa 0.4.6^108^ (*ϕ* = 0.933, *p*-value = 0.660, two-sided). Simple Poisson regressions fitted under the same protocol systematically showed severe under-dispersion in their residuals (results not shown).

### Embryonic development *in vivo*

We observed m1 cusp development *in vivo* at intervals of 24 hours, from embryonic developmental day 14.5 (E14.5) to embryonic developmental day 17 (E17) in mouse embryos (*Mus musculus*) and to E18 in bank vole embryos (*Clethrionomys glareolus*). The appearance of the vaginal plug was taken as the developmental stage 0 (E0). Developmental stages were defined using limb morphology and tooth comparison. Pregnant females were euthanised by CO_2_ gas inhalation and neck dislocation before harvesting embryos (one individual per stage; see Ethics declarations below). Dissected half lower jaws were fixed overnight in 4% paraformaldehyde (PFA), then dehydrated in an overnight series of 50% to 70% ethanol. Samples were stained overnight with 0.3% (weight/volume) phosphotungstic acid (PTA) in 70% ethanol to increase soft tissue contrast for X-ray computed microtomography (microCT) scanning and stored at 4°C in 70% ethanol before scanning^114^. We scanned samples in 70% ethanol using a custom-built microCT system Nanotom 180 NF (with an effective voxel size between 4 and 8 μm). We used ImageJ 1.45c^83^ and AMIRA ® 5.5.0^115^ for 3D segmenting the developing tooth epithelium and mesenchyme. 3D renders of mouse and bank vole m1 development were adapted from Renvoisé *et al.*^37^

### Tooth explant cultures *in vitro*

First lower molar tooth germs were dissected from embryos of pregnant *Clethrionomys glareolus* females (see Ethics declarations below) at embryonic days E13 (*n* = 9) and E14 (*n* = 9) (see above) for six to seven days of *in vitro* culture in a Trowell-type organ culture system^116^. The culture medium (DMEM) was changed every second day and supplemented 1:1 with F12 (Ham’s Nutrient Mix: Life Technologies) and with 100 ng/mL ascorbic acid (Merck) from the second day onwards. Bright-field pictures were taken every 24 hours with an Olympus SZX9 microscope.

### Statistical information

All analyses and statistical tests were replicated twice independently with consistent results unless specified otherwise (five independent replicates for the null and variable rates models of evolution of molar dimensions).

## Ethics declarations

All aspects of the study involving animals (housing, care, captive breeding, procedures, experiments) were approved by the Laboratory Animal Centre (LAC) of the University of Helsinki and the National Animal Experiment Board (ELLA) of Finland under the license number KEK16-021.

## Data availability statement

All datasets generated and analysed during the current study (tip-state dataset, phylogenies, *in vivo* and *in vitro* measurements) are available as Supplementary Information. CT-scan data are available upon request and will be made available on a public repository upon publication.

## Supporting information

Supplementary Information guide

## Acknowledgements

This work used the services of the University of Helsinki X-Ray Micro-CT Laboratory, funded also by the Helsinki Institute of Life Science (HiLIFE) under the HAIP platform. The authors wish to thank Heikki Suhonen and Aki Kallonen for access to the facility and assistance in scanning specimens, Jean-Pierre Quéré, and Jukka Jernvall, Mikael Fortelius, and the Helsinki Evo-Devo community for discussions. This work was supported by funds from the Center for International Mobility scholarship program (CIMO; to F.L.), the Integrative Life Science doctoral program (ILS; to F.L.), and the Academy of Finland (to É.R.).

## Author contributions

G.E., É.R., and F.L. designed the study. É.R. acquired the embryo samples, imaged them, performed the *in vitro* experiments, and collected the data from the embryos. F.L. and É.R. compiled the fossil and extant taxa dataset, and F.L. character-coded the dataset. F.L. assembled the informal super-tree. F.L. and J.C. analysed the data. F.L., É.R., and I.J.C. interpreted the results. F.L. wrote the first draft and produced the figures and supplementary information. All authors took part in the review and editing process and approved the final version of the manuscript.

## Additional information

Supplementary information is available for this paper. Correspondence and requests for materials should be addressed to F.L.

---

**Extended Data Fig. 1.**
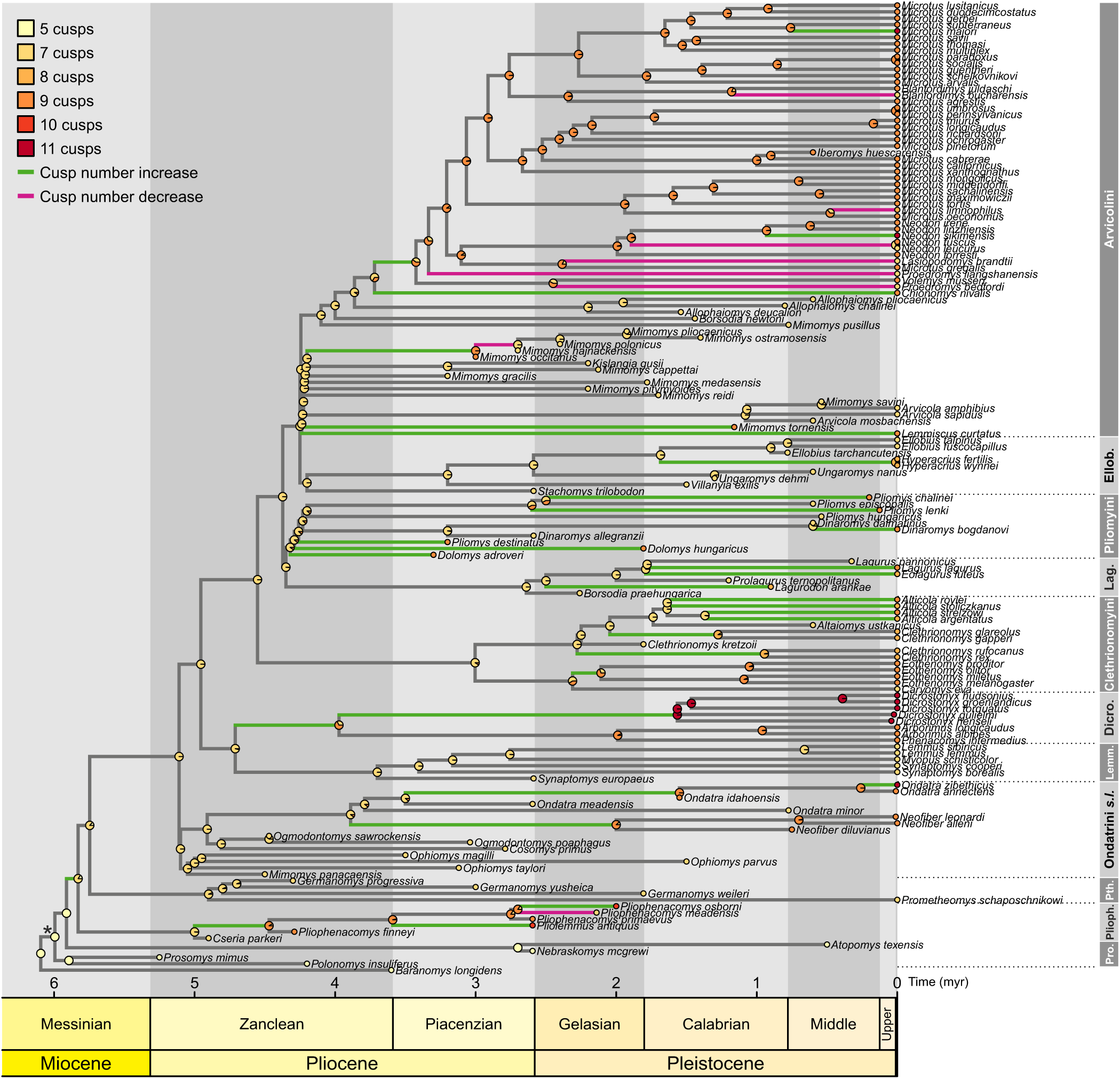
Variations in first lower molar cusp number throughout the evolution of Arvicolinae. Observed first lower molar cusp numbers (tips) and maximum likelihood ancestral state estimations at nodes (pie charts) across a phylogeny of 151 species of Arvicolinae under an extended M*k* model (see Methods). Branch colours denote cusp number increases (green), decreases (magenta), or stasis (grey). Pro.: grade ‘Promimomyini’. Plioph.: Pliophenacomyini. Pth.: Prometheomyini. Ondatrini *s.l.*: Ondatrini *sensu lato* (including *Ondatra*, *Neofiber*, and related stem-taxa). Lemm.: Lemmini. Dicro.: Dicrostonychini *sensu lato* (including *Dicrostonyx*, *Arborimus*, and *Phenacomys*). Lag.: Lagurini. Ellob.: Ellobiusini. Asterisk: most recent common ancestor (MRCA) of Arvicolinae.

**Extended Data Fig. 2.**
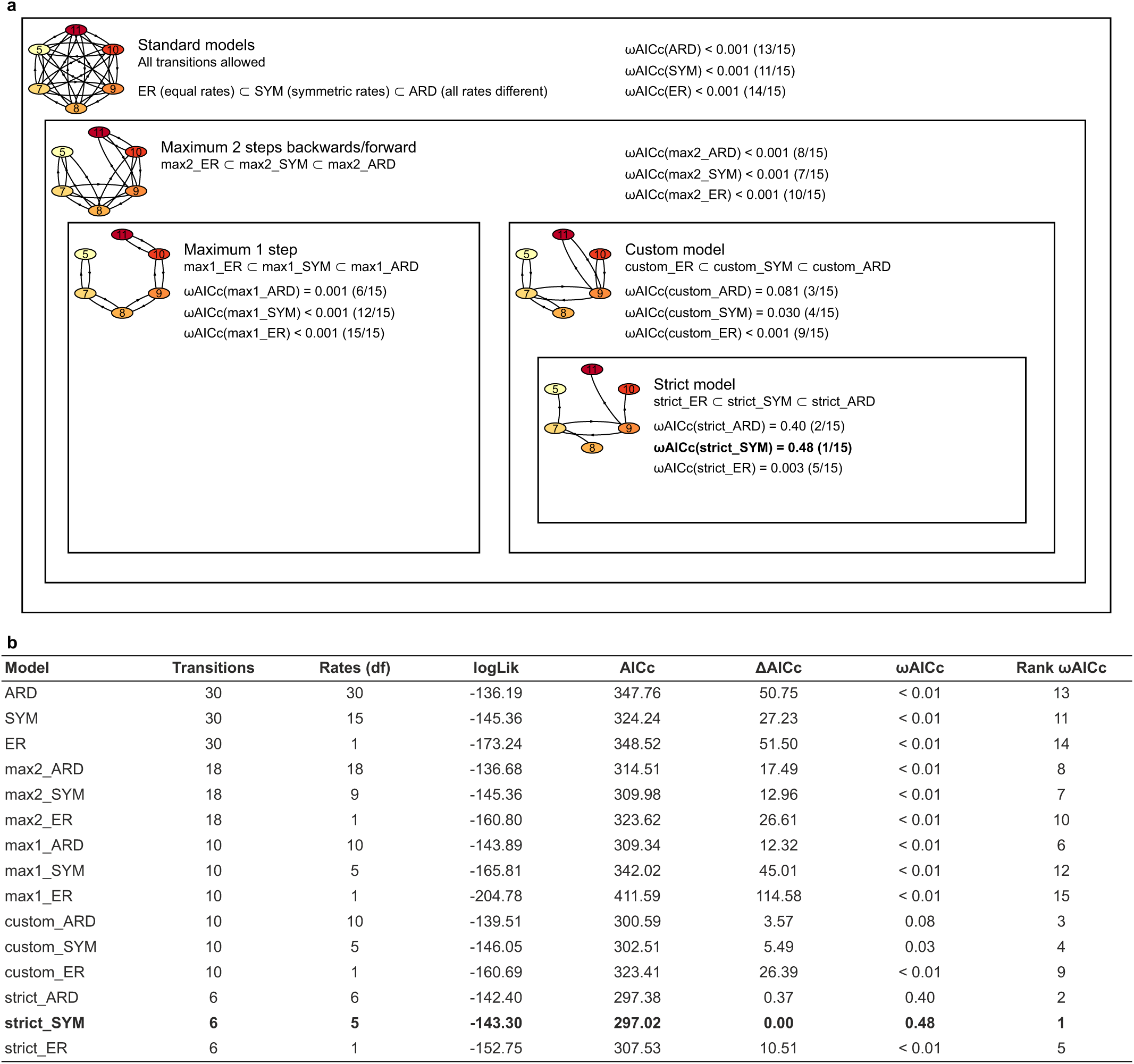
Models of cusp number evolution tested in the study. **a**, Schematic representation of the candidate set of nested M*k* models of cusp number evolution fitted to our phylogenetic dataset of first lower molar cusp counts in Arvicolinae, with arrows representing the character transitions allowed in each model category. Models with similar topologies (*i.e.*, character transitions) but distinct rates of transitions are listed in the corresponding boxes, along with their relative statistical support (*ω*AICc) and rank. *ω*AICc: weighed corrected Akaike Information Criterion. **b**, Relative statistical support for each M*k* model of cusp number evolution fitted to our evolutionary dataset of first lower molar cusp counts in Arvicolinae based on the corrected Akaike Information Criterion (AICc). The best-supported model (*i.e.*, with the highest AICc weight) is highlighted in bold. *ω*AICc: AICc weight. ΔAICc: AICc differential to the best model’s AICc value. df: degrees of freedom. logLik: marginal log-likelihood. ARD: all rates are different. SYM: symmetric rates (forces rate[A➡B] = rate[B➡A]). ER: equal rates.

**Extended Data Fig. 3.**
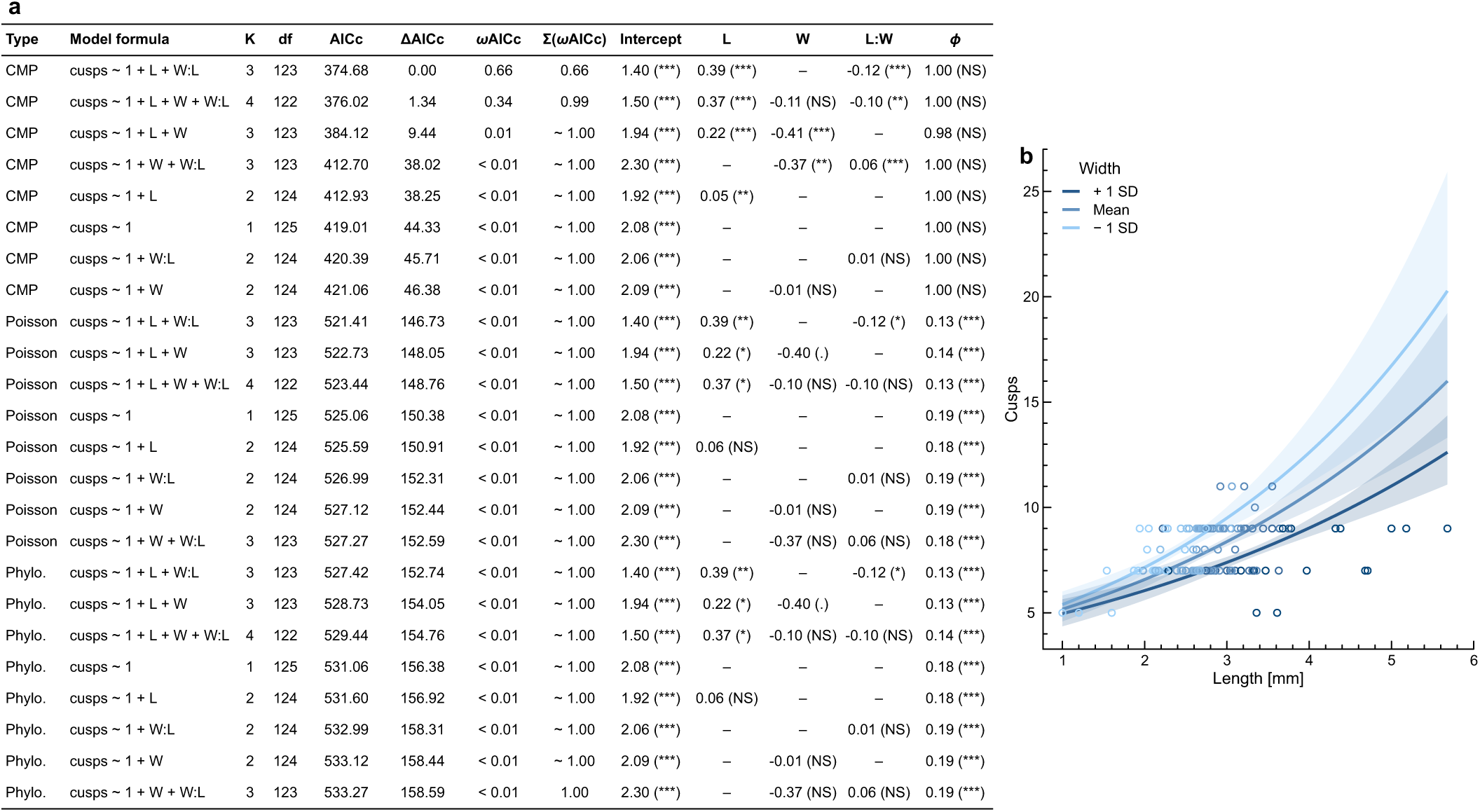
Longer first lower molars bear more cusps throughout the evolution of Arvicolinae, though broader teeth weaken the effect of length on cusp number. **a**, Conway–Maxwell–Poisson (CMP), Poisson, and phylogenetic Poisson (Phylo.) regression models fitted to our evolutionary dataset of first lower molar cusp counts and dimensions in Arvicolinae (*n* = 127) ranked by decreasing relative statistical support based on the corrected Akaike Information Criterion (AICc). K: number of model parameters. df: degrees of freedom. ΔAICc: AICc differential to the best model AICc value. *ω*AICc: AICc weight. Ʃ(*ω*AICc): cumulated sum of AICc weights from best to least supported model. L: first lower molar length. W: first lower molar width. L:W: interaction term between first lower molar width and length. *ϕ*: dispersion. NS: *p*-value > 0.05. (.): 0.01 < *p* < 0.05. (*): 0.001 < *p* < 0.01. (**): 0.0001 < *p* < 0.001. (***): *p* < 0.0001. See Extended Data Table 1 for the full statistical reporting of parameter estimates. **b**, First lower molar cusp number as a function of length under the best-supported model (cusps ∼ 1 + L + W:L, CMP regression) with regression slopes and their 95% confidence intervals (shaded areas) under three values of width (average value and average value ±1 standard deviation). The colours of the data points denote width values. See Methods for the full statistical reporting of model diagnostics.

**Extended Data Fig. 4.**
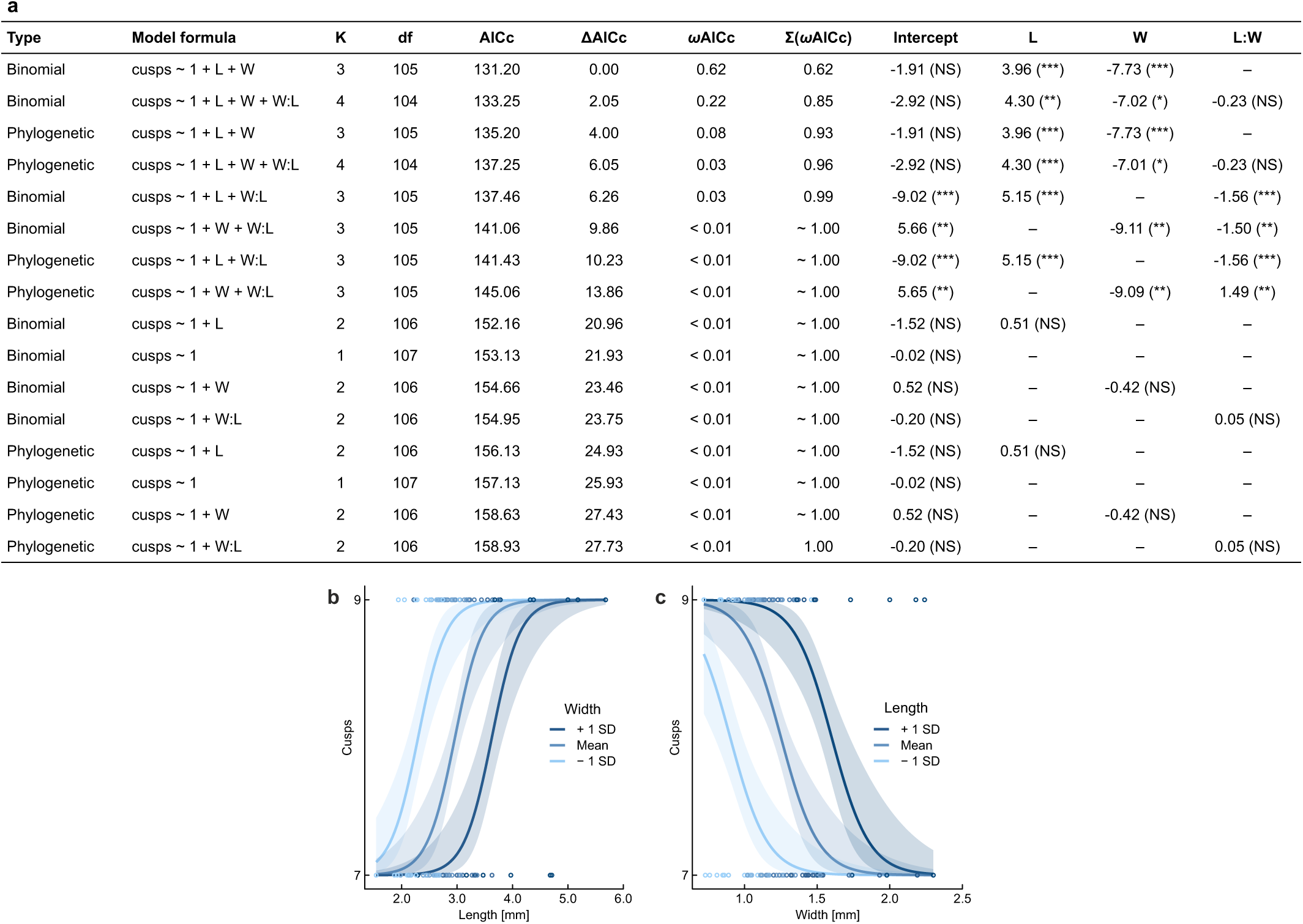
Longer, narrower teeth characterise the transition from seven to nine cusps in the evolution of the first lower molar of Arvicolinae. **a**, Phylogenetic and non-phylogenetic logistic binomial regression models fitted to our dataset of cusp counts and dimensions limited to species bearing seven- or nine-cusped first lower molars (*n* = 109) ranked by decreasing relative support based on the corrected Akaike Information Criterion (AICc). K: number of parameters. df: degrees of freedom. ΔAICc: AICc differential to the best model. *ω*AICc: AICc weight. Ʃ(*ω*AICc): cumulated sum of AICc weights from best to least supported model. L: m1 length. W: m1 width. L:W: interaction term between m1 width and length. NS: *p*-value > 0.05. (.): 0.01 < *p* < 0.05. (*): 0.001 < *p* < 0.01. (**): 0.0001 < *p* < 0.001. (***): *p* < 0.0001. See Extended Data Table 1 for the full statistical reporting of parameter estimates. **b– c**, m1 cusp number as a binary state function (seven or nine cusps) of length **(b)** and width **(c)** under the best-supported model (cusps ∼ 1 + L + W, non-phylogenetic logistic binomial regression) with regression slopes and their 95% confidence intervals represented under three values of width and length, respectively (average value and average value ±1 standard deviation). Data point colour denotes width and length values, respectively. See Methods for model diagnostics.

**Extended Data Fig. 5.**
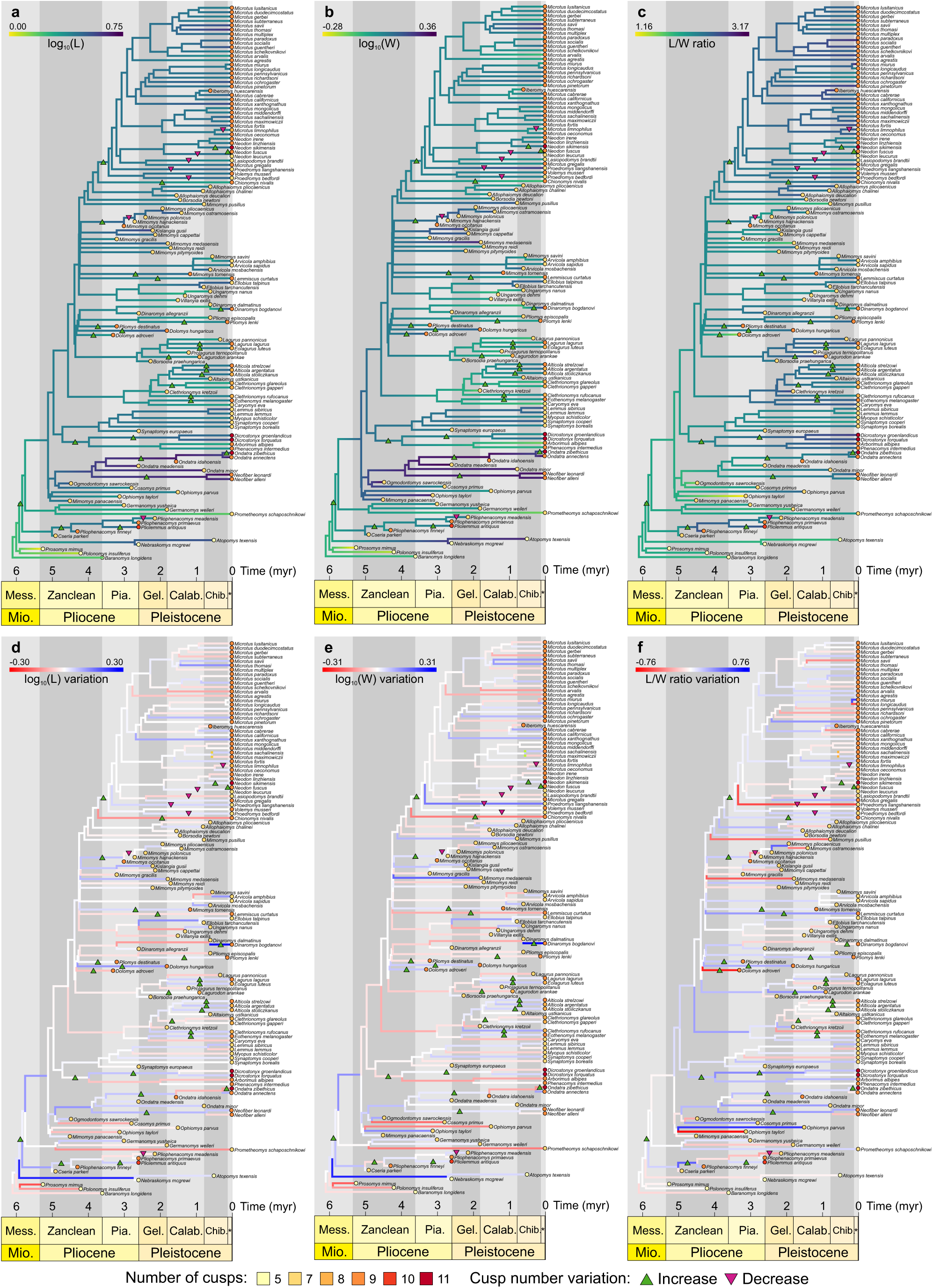
Evolution of first lower molar dimensions in Arvicolinae. **a–c**, Reconstructed evolution of log-transformed length (log_10_(L)) **(a)**, log-transformed width (log_10_(W)) **(b)**, and length/width ratio (L/W ratio) **(c)** of arvicoline first lower molars under a Brownian Motion (BM) process across a phylogeny of 126 species of Arvicolinae and one outgroup taxon (*Baranomys longidens*). **d–f**, Implied changes in the log-transformed length (log_10_(L)) **(d)**, log-transformed width (log_10_(W)) **(e)**, and length/width ratio (L/W ratio) **(f)** of the first lower molar. Triangles indicate lineages with increasing (green) or decreasing cusp number (magenta) based on our maximum likelihood ancestral reconstructions of m1 cusp number (see Extended Data Fig. 9). Mess.: Messinian. Pia.: Piacenzian. Gel.: Gelasian. Calab.: Calabrian. Chib.: Chibanian. Asterisk: Upper Pleistocene.

**Extended Data Fig. 6.**
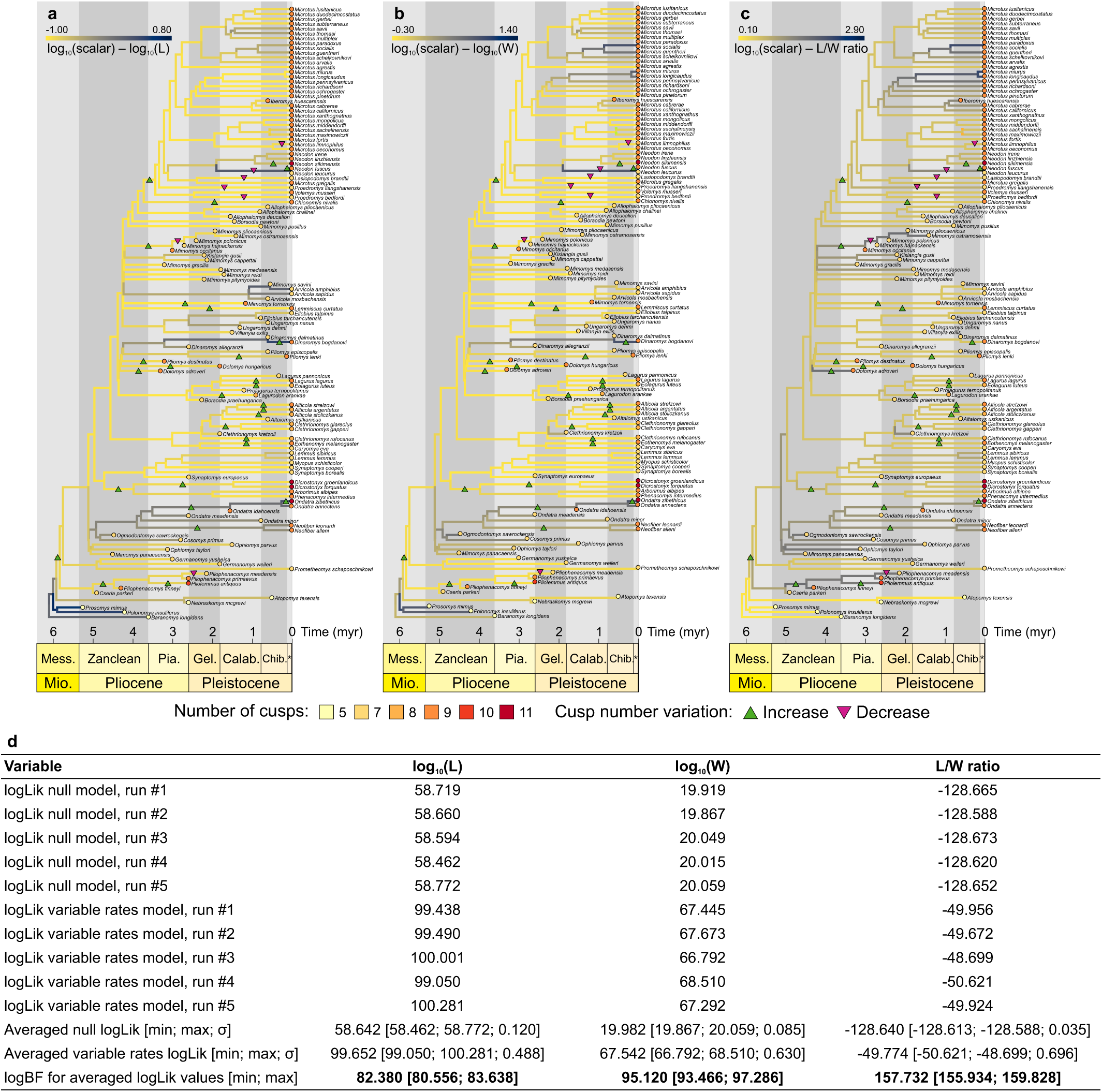
Rates of evolution of first lower molar dimensions in Arvicolinae. **a-c**, Rates of evolution of the log-transformed length (log_10_(L)) **(a)**, log-transformed width (log_10_(W)) **(b)**, and length/width ratio (L/W ratio) **(c)** of the first lower molar in 126 Arvicolinae based on ancestral state reconstructions under a heterogeneous “variable rates” model of evolution (allowing local heterogeneity in the character transition rates). Triangles indicate lineages with increasing (green) or decreasing cusp number (magenta) based on our maximum likelihood ancestral reconstructions of first lower molar cusp number (see Extended Data Fig. 9). Mess.: Messinian. Pia.: Piacenzian. Gel.: Gelasian. Calab.: Calabrian. Chib.: Chibanian. Asterisk: Upper Pleistocene. **d**, Marginal log-likelihoods (logLik) for five independent runs of constant and “variable rates” models of trait evolution of three first lower molar linear measurements (log_10_-transformed length (log_10_(L)), log_10_-transformed width (log_10_(W), and length/width ratio), with their associated log Bayes Factor (logBF). Values in bold (logBF > 10) indicate very strong statistical support for the more complex model (*i.e.*, the “variable rates” model).

**Extended Data Fig. 7.**
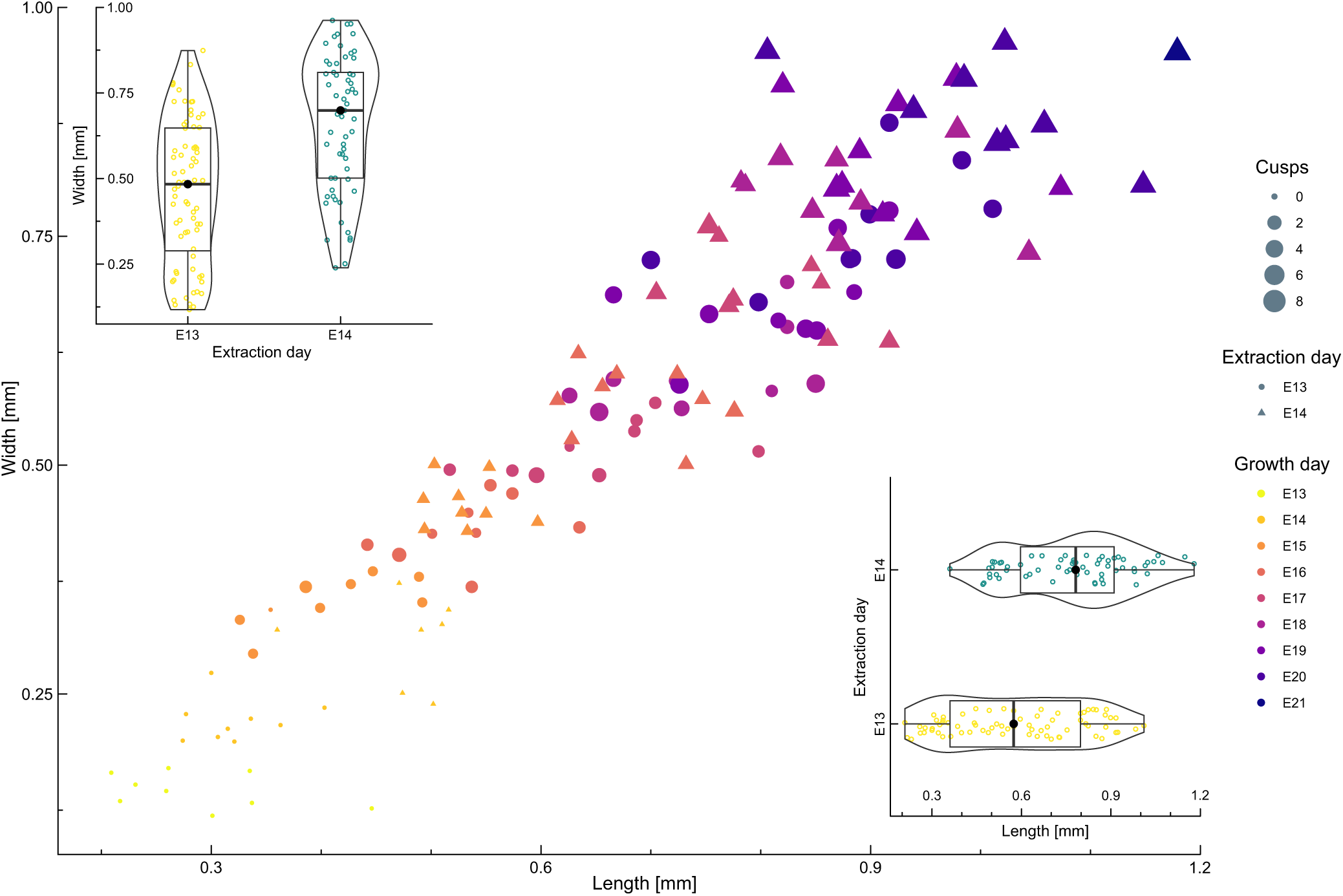
First lower molar germs of bank voles extracted earlier for culture grow into smaller, less cuspidate teeth *in vitro*. Cusp counts and linear measurements for bank vole first lower molar germs developing *in vitro* after extraction at developmental stage E13 (circles) or E14 (triangles) (*n* = 133). Data point size indicates cusp number and colour, the equivalent developmental stage. Insets: length (bottom right) and width (top left) of the cultured germs extracted at developmental stage E13 (*n* = 72) or E14 (*n* = 61) pooled over the entire duration of the explant culture. Boxes contain 50% of the data points (25^th^ to 75^th^ percentile). Thick black lines and black dots indicate the median and mean for each experimental condition, respectively. Whiskers span the entire range of the data (minimum to maximum value; no outliers were detected). Violin plots illustrate data point density.

**Extended Data Fig. 8.**
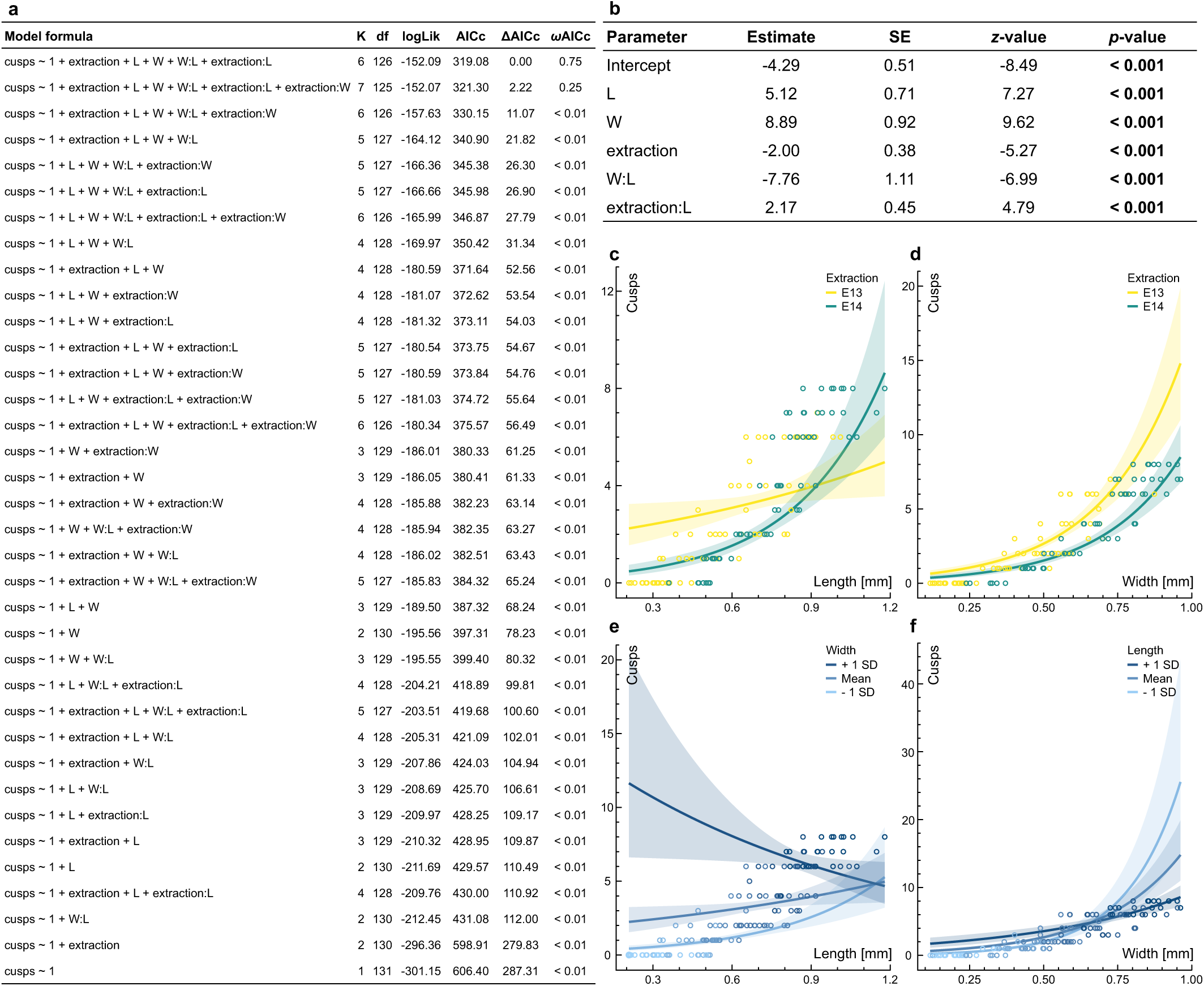
Effects of germ dimensions and extraction day on cusp number in first lower molars of bank voles developing *in vitro*. **a**, CMP regression models fitted to the dataset of molar complexity and dimensions *in vitro* (*n* = 133) ranked by relative statistical support based on the corrected Akaike Information Criterion (AICc). K: number of model parameters. df: degrees of freedom. logLik: log-likelihood. ΔAICc: AICc differential to the best model’s AICc value. *ω*AICc: AICc weight. L: first lower molar (m1) length. W: m1 width. extraction: day of extraction. W:L: interaction term between m1 width and length. extraction:L: interaction term between extraction day and m1 length. extraction:W: interaction term between extraction day and m1 width. **b**, Parameter estimates for the best-supported Conway–Maxwell– Poisson model (cusps ∼ 1 + L + W + extraction + W:L + extraction:L). SE: standard error. Significant (*p* < 0.05) relationships between predictors and response are in bold. **c–f**, Cusp number as a function of length **(c, e)** and width **(d, f)** for m1 germs developing *in vitro* (*n* = 133) after extraction at developmental stage E13 (yellow, *n* = 72) and E14 (green, *n* = 61) under the best-supported CMP fit. Regression slopes and their 95% confidence interval are represented for each experimental condition **(c, d)** and for the pooled *in vitro* dataset under three values of width **(e)** and length **(f)** (average value and average value ±1 standard deviation), with the colours of the data points denoting width and length values, respectively. See Methods for the full statistical reporting.

**Extended Data Fig. 9.**
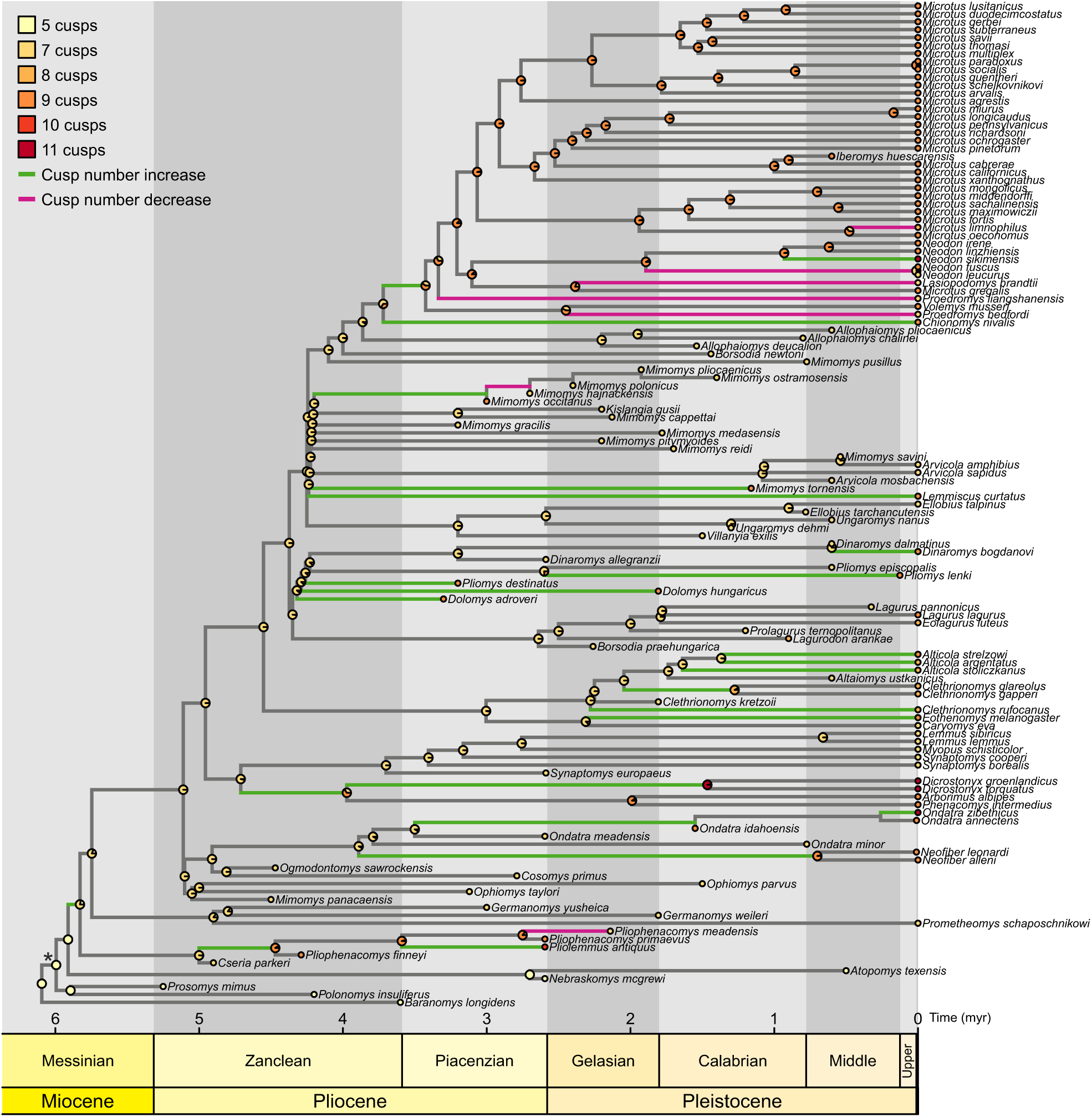
Variations in first lower molar cusp number throughout the evolution of Arvicolinae, limited to species with known molar dimensions. Observed first lower molar cusp numbers (tips) and maximum likelihood ancestral state estimations at nodes (pie charts) across a phylogeny of 126 species of Arvicolinae with known molar dimensions under an extended M*k* model (see Methods). Branch colours denote cusp number increases (green), decreases (magenta), or stasis (grey). Asterisk: most recent common ancestor (MRCA) of Arvicolinae.

**Extended Data Table 1.** Parameter estimates for the different regression models fitted to the evolutionary dataset of m1 cusp number. Phylogenetic and non-phylogenetic regression models fitted to the full evolutionary dataset of first lower molar cusp counts and dimensions in Arvicolinae (*n* = 127) and its subset limited to species with a seven- or nine-cusped m1 (*n* = 109). CMP: Conway–Maxwell–Poisson regression. Poisson (phy): phylogenetic Poisson regression. GEE: phylogenetic Poisson regression fitted with the method of Generalised Estimating Equations^109^. Binomial (phy): phylogenetic logistic regression. IG10: phylogenetic logistic regression fitted with the method of Ives & Garland^111^. L: first lower molar length. W: first lower molar width. W:L: interaction term between first lower molar width and length. SE: standard error. Statistically significant (*p* < 0.05) coefficients are in bold.

| Regression | Model formula | Intercept | SE | p | L | SE | p | W | SE | p | L:W | SE | p |
| --- | --- | --- | --- | --- | --- | --- | --- | --- | --- | --- | --- | --- | --- |
| CMP | m1 ~ 1 | 2.08 | 0.01 | < 0.001 | – | – | – | – | – | – | – | – | – |
| Poisson | m1 ~ 1 | 2.08 | 0.03 | < 0.001 | – | – | – | – | – | – | – | – | – |
| Poisson (phy) | m1 ~ 1 | 2.08 | 0.03 | < 0.001 | – | – | – | – | – | – | – | – | – |
| GEE | m1 ~ 1 | 1.98 | 0.06 | < 0.001 | – | – | – | – | – | – | – | – | – |
| Binomial | m1 ~ 1 | -0.02 | 0.19 | 0.924 | – | – | – | – | – | – | – | – | – |
| Binomial (phy) | m1 ~ 1 | -0.02 | 0.19 | 0.924 | – | – | – | – | – | – | – | – | – |
| IG10 | m1 ~ 1 | -0.03 | 0.20 | 0.865 | – | – | – | – | – | – | – | – | – |
| CMP | m1 ~ 1 + L | 1.92 | 0.06 | < 0.001 | 0.05 | 0.02 | 0.003 | – | – | – | – | – | – |
| Poisson | m1 ~ 1 + L | 1.92 | 0.13 | < 0.001 | 0.06 | 0.04 | 0.213 | – | – | – | – | – | – |
| Poisson (phy) | m1 ~ 1 + L | 1.92 | 0.13 | < 0.001 | 0.06 | 0.04 | 0.213 | – | – | – | – | – | – |
| GEE | m1 ~ 1 + L | 2.4 | 0.08 | < 0.001 | -0.16 | 0.02 | < 0.001 | – | – | – | – | – | – |
| Binomial | m1 ~ 1 + L | -1.52 | 0.91 | 0.096 | 0.51 | 0.31 | 0.094 | – | – | – | – | – | – |
| Binomial (phy) | m1 ~ 1 + L | -1.52 | 0.92 | 0.100 | 0.51 | 0.31 | 0.096 | – | – | – | – | – | – |
| IG10 | m1 ~ 1 + L | -1.20 | 1.06 | 0.257 | 0.41 | 0.28 | 0.147 | – | – | – | – | – | – |
| CMP | m1 ~ 1 + W | 2.09 | 0.05 | < 0.001 | – | – | – | -0.01 | 0.04 | 0.822 | – | – | – |
| Poisson | m1 ~ 1 + W | 2.09 | 0.12 | < 0.001 | – | – | – | -0.01 | 0.09 | 0.921 | – | – | – |
| Poisson (phy) | m1 ~ 1 + W | 2.09 | 0.12 | < 0.001 | – | – | – | -0.01 | 0.09 | 0.921 | – | – | – |
| GEE | m1 ~ 1 + W | 2.29 | 0.07 | < 0.001 | – | – | – | -0.27 | 0.03 | < 0.001 | – | – | – |
| Binomial | m1 ~ 1 + W | 0.52 | 0.75 | 0.489 | – | – | – | -0.42 | 0.57 | 0.459 | – | – | – |
| Binomial (phy) | m1 ~ 1 + W | 0.52 | 0.76 | 0.493 | – | – | – | -0.42 | 0.58 | 0.463 | – | – | – |
| IG10 | m1 ~ 1 + W | 0.55 | 0.82 | 0.502 | – | – | – | -0.48 | 0.54 | 0.375 | – | – | – |
| CMP | m1 ~ 1 + W:L | 2.06 | 0.03 | < 0.001 | – | – | – | – | – | – | 0.01 | 0.01 | 0.395 |
| Poisson | m1 ~ 1 + W:L | 2.06 | 0.07 | < 0.001 | – | – | – | – | – | – | 0.01 | 0.02 | 0.710 |
| Poisson (phy) | m1 ~ 1 + W:L | 2.06 | 0.07 | < 0.001 | – | – | – | – | – | – | 0.01 | 0.02 | 0.710 |
| GEE | m1 ~ 1 + W:L | 2.16 | 0.07 | < 0.001 | – | – | – | – | – | – | -0.05 | 0.01 | < 0.001 |
| Binomial | m1 ~ 1 + W:L | -0.20 | 0.41 | 0.621 | – | – | – | – | – | – | 0.05 | 0.09 | 0.612 |
| Binomial (phy) | m1 ~ 1 + W:L | -0.20 | 0.42 | 0.624 | – | – | – | – | – | – | 0.05 | 0.09 | 0.615 |
| IG10 | m1 ~ 1 + W:L | -0.78 | 0.62 | 0.206 | – | – | – | – | – | – | 0.08 | 0.09 | 0.419 |
| CMP | m1 ~ 1 + L + W | 1.94 | 0.05 | < 0.001 | 0.22 | 0.03 | < 0.001 | -0.41 | 0.07 | < 0.001 | – | – | – |
| Poisson | m1 ~ 1 + L + W | 1.94 | 0.13 | < 0.001 | 0.22 | 0.09 | 0.012 | -0.40 | 0.19 | 0.030 | – | – | – |
| Poisson (phy) | m1 ~ 1 + L + W | 1.94 | 0.13 | < 0.001 | 0.22 | 0.09 | 0.012 | -0.40 | 0.19 | 0.029 | – | – | – |
| GEE | m1 ~ 1 + L + W | 2.36 | 0.08 | < 0.001 | -0.06 | 0.03 | 0.04 | -0.18 | 0.05 | < 0.001 | – | – | – |
| Binomial | m1 ~ 1 + L + W | -1.91 | 1.05 | 0.070 | 3.96 | 1.00 | < 0.001 | -7.73 | 2.05 | < 0.001 | – | – | – |
| Binomial (phy) | m1 ~ 1 + L + W | -1.91 | 1.05 | 0.070 | 3.96 | 1.00 | < 0.001 | -7.73 | 2.05 | < 0.001 | – | – | – |
| IG10 | m1 ~ 1 + L + W | -1.48 | 1.16 | 0.200 | 2.98 | 0.83 | < 0.001 | -5.95 | 1.73 | < 0.001 | – | – | – |
| CMP | m1 ~ 1 + L + W:L | 1.40 | 0.09 | < 0.001 | 0.39 | 0.05 | < 0.001 | – | – | – | -0.12 | 0.02 | < 0.001 |
| Poisson | m1 ~ 1 + L + W:L | 1.40 | 0.25 | < 0.001 | 0.39 | 0.14 | 0.007 | – | – | – | -0.12 | 0.05 | 0.013 |
| Poisson (phy) | m1 ~ 1 + L + W:L | 1.40 | 0.25 | < 0.001 | 0.39 | 0.14 | 0.006 | – | – | – | -0.12 | 0.05 | 0.013 |
| GEE | m1 ~ 1 + L + W:L | 2.16 | 0.11 | < 0.001 | -0.01 | 0.05 | 0.921 | – | – | – | -0.05 | 0.02 | 0.002 |
| Binomial | m1 ~ 1 + L + W:L | -9.02 | 2.41 | < 0.001 | 5.15 | 1.38 | < 0.001 | – | – | – | -1.56 | 0.45 | < 0.001 |
| Binomial (phy) | m1 ~ 1 + L + W:L | -9.02 | 2.43 | < 0.001 | 5.15 | 1.39 | < 0.001 | – | – | – | -1.56 | 0.45 | < 0.001 |
| IG10 | m1 ~ 1 + L + W:L | -8.51 | 2.39 | < 0.001 | 4.84 | 1.32 | < 0.001 | – | – | – | -1.50 | 0.44 | < 0.001 |
| CMP | m1 ~ 1 + W + W:L | 2.30 | 0.08 | < 0.001 | – | – | – | -0.37 | 0.12 | 0.001 | 0.06 | 0.02 | < 0.001 |
| Poisson | m1 ~ 1 + W + W:L | 2.30 | 0.19 | < 0.001 | – | – | – | -0.37 | 0.28 | 0.181 | 0.06 | 0.05 | 0.162 |
| Poisson (phy) | m1 ~ 1 + W + W:L | 2.30 | 0.19 | < 0.001 | – | – | – | -0.37 | 0.28 | 0.181 | 0.06 | 0.05 | 0.162 |
| GEE | m1 ~ 1 + W + W:L | 2.22 | 0.08 | < 0.001 | – | – | – | -0.11 | 0.09 | 0.225 | -0.03 | 0.02 | 0.061 |
| Binomial | m1 ~ 1 + W + W:L | 5.66 | 1.81 | 0.002 | – | – | – | -9.11 | 2.80 | 0.001 | -1.50 | 0.48 | 0.002 |
| Binomial (phy) | m1 ~ 1 + W + W:L | 5.65 | 1.82 | 0.002 | – | – | – | -9.09 | 2.81 | 0.001 | 1.49 | 0.48 | 0.002 |
| IG10 | m1 ~ 1 + W + W:L | 7.42 | 2.05 | < 0.001 | – | – | – | -13.04 | 3.42 | < 0.001 | 2.22 | 0.60 | < 0.001 |
| CMP | m1 ~ 1 + L + W + W:L | 1.50 | 0.14 | < 0.001 | 0.37 | 0.06 | < 0.001 | -0.11 | 0.11 | 0.360 | -0.10 | 0.03 | 0.001 |
| Poisson | m1 ~ 1 + L + W + W:L | 1.50 | 0.39 | < 0.001 | 0.37 | 0.16 | 0.017 | -0.10 | 0.32 | 0.747 | -0.10 | 0.08 | 0.241 |
| Poisson (phy) | m1 ~ 1 + L + W + W:L | 1.50 | 0.39 | < 0.001 | 0.37 | 0.16 | 0.017 | -0.10 | 0.32 | 0.747 | -0.10 | 0.08 | 0.241 |
| GEE | m1 ~ 1 + L + W + W:L | 2.33 | 0.16 | < 0.001 | -0.05 | 0.06 | 0.387 | -0.16 | 0.11 | 0.155 | -0.01 | 0.03 | 0.841 |
| Binomial | m1 ~ 1 + L + W + W:L | -2.92 | 3.37 | 0.386 | 4.30 | 1.47 | 0.003 | -7.02 | 3.01 | 0.020 | -0.23 | 0.71 | 0.749 |
| Binomial (phy) | m1 ~ 1 + L + W + W:L | -2.92 | 3.37 | 0.386 | 4.30 | 1.47 | 0.004 | -7.01 | 3.01 | 0.020 | -0.23 | 0.71 | 0.749 |
| IG10 | m1 ~ 1 + L + W + W:L | -2.94 | 3.18 | 0.354 | 3.83 | 1.42 | 0.007 | -5.63 | 2.69 | 0.036 | -0.35 | 0.71 | 0.623 |

## References

1 Darwin, C. On the Origin of Species by Means of Natural Selection or the Preservation of the Favoured Races in the Struggle for Life. (John Murray, Albemarle Street, London, 1859).

2 Alberch, P. Ontogenesis and morphological diversification. Am. Zool. 20, 653–667 (1980).

3 Kimura, M. The Neutral Theory of Molecular Evolution. (Cambridge University Press, 1985).

4 Erwin, D. H. Developmental push or environmental pull? The causes of macroevolutionary dynamics. Hist. Philos. Life Sci. 39, 36 (2017).

5 Hunter, J. P. & Jernvall, J. The hypocone as a key innovation in mammalian evolution. Proc. Natl. Acad. Sci. USA 92, 10718–10722 (1995).

6 Ungar, P. S. Mammal Teeth: Origin, Evolution, and Diversity. (JHU Press, 2010).

7 Couzens, A. M., Sears, K. E. & Rücklin, M. Developmental influence on evolutionary rates and the origin of placental mammal tooth complexity. Proc. Natl. Acad. Sci. USA 118, e2019294118 (2021).

8 Jernvall, J., Keränen, S. V. & Thesleff, I. Evolutionary modification of development in mammalian teeth: quantifying gene expression patterns and topography. Proc. Natl. Acad. Sci. USA 97, 14444–14448 (2000).

9 Salazar-Ciudad, I. & Jernvall, J. A computational model of teeth and the developmental origins of morphological variation. Nature 464, 583–586 (2010).

10 Jablonski, D. Approaches to macroevolution: 1. General concepts and origin of variation. Evol. Biol. 44, 427–450 (2017).

11 Simpson, G. G. Tempo and Mode in Evolution. (Columbia University Press, 1944).

12 Stroud, J. T. & Losos, J. B. Ecological opportunity and adaptive radiation. Annu. Rev. Ecol., Evol. Syst. 47, 507–532 (2016).

13 Gould, S. J. Ontogeny and Phylogeny. (Harvard University Press, 1977).

14 Gould, S. J. & Lewontin, R. C. The spandrels of San Marco and the Panglossian paradigm: a critique of the adaptationist programme. *Proc. R. Soc. Lond.*, Ser. B: Biol. Sci. 205, 79 (1979).

15 Goldschmidt, R. B. The Material Basis of Evolution. Vol. 28 (Yale University Press, 1940).

16 Godfrey-Smith, P. Philosophy of Biology. (Princeton University Press, 2014).

17 Carroll, S. B. Evo-devo and an expanding evolutionary synthesis: a genetic theory of morphological evolution. Cell 134, 25–36 (2008).

18 Lowe, C. B. et al. Three periods of regulatory innovation during vertebrate evolution. Science 333, 1019–1024 (2011).

19 Maynard Smith, J., et al. Developmental constraints and evolution. Q. Rev. Biol. 60, 265–287 (1985).

20 Beldade, P., Koops, K. & Brakefield, P. M. Developmental constraints versus flexibility in morphological evolution. Nature 416, 844–847 (2002).

21 Alberch, P., Gould, S. J., Oster, G. F. & Wake, D. B. Size and shape in ontogeny and phylogeny. Paleobiology 5, 296–317 (1979).

22 Escarguel, G., Fara, E., Brayard, A. & Legendre, S. Biodiversity is not (and never has been) a bed of roses! *C. R*. Biol. 334, 351–359 (2011).

23 Duran-Nebreda, S. et al. On the multiscale dynamics of punctuated evolution. Trends Ecol. Evol. (2024).

24 Jablonski, D. Developmental bias, macroevolution, and the fossil record. Evol. Dev. 22, 103–125 (2020).

25 Jernvall, J., Kettunen, P., Karavanova, I., Martin, L. B. & Thesleff, I. Evidence for the role of the enamel knot as a control center in mammalian tooth cusp formation: non-dividing cells express growth stimulating Fgf-4 gene. Int. J. Dev. Biol. 38, 463–469 (1994).

26 Vaahtokari, A., Åberg, T., Jernvall, J., Keränen, S. & Thesleff, I. The enamel knot as a signaling center in the developing mouse tooth. Mech. Dev. 54, 39–43 (1996).

27 Salazar-Ciudad, I. & Jernvall, J. A gene network model accounting for development and evolution of mammalian teeth. Proc. Natl. Acad. Sci. USA 99, 8116–8120 (2002).

28 Harjunmaa, E. et al. On the difficulty of increasing dental complexity. Nature 483, 324–327 (2012).

29 Harjunmaa, E. et al. Replaying evolutionary transitions from the dental fossil record. Nature 512, 44–48 (2014).

30 Burroughs, R. W. Phylogenetic and developmental constraints dictate the number of cusps on molars in rodents. Sci. Rep. 9, 10902 (2019).

31 Chaline, J., Brunet-Lecomte, P., Montuire, S. & Viriot, L. Anatomy of the arvicoline radiation (Rodentia): palaeogeographical, palaeoecological history and evolutionary data. Ann. Zool. Fenn. 36, 239–267 (1999).

32 Steppan, S. J. & Schenk, J. J. Muroid rodent phylogenetics: 900-species tree reveals increasing diversification rates. PLoS One 12, e0183070 (2017).

33 Renvoisé, É. et al. Evolution of mammal tooth patterns: New insights from a developmental prediction model. Evolution 63, 1327–1340 (2009).

34 Cantalapiedra, J. L. et al. The rise and fall of proboscidean ecological diversity. *Nat*. Ecol. Evol. 5, 1266–1272 (2021).

35 Fejfar, O., Heinrich, W.-D., Kordos, L. & Maul, L. C. Microtoid cricetids and the early history of arvicolids (Mammalia, Rodentia). Palaeontol. Electron. 14, 1–38 (2011).

36 Christensen, M. M. et al. The developmental basis for scaling of mammalian tooth size. Proc. Natl. Acad. Sci. USA 120, e2300374120 (2023).

37 Renvoisé, É. et al. Mechanical constraint from growing jaw facilitates mammalian dental diversity. Proc. Natl. Acad. Sci. USA 114, 9403–9408 (2017).

38 Renvoisé, É. & Montuire, S. in Evolution of the Rodents: Advances in Phylogeny, Functional Morphology and Development (eds P. G. Cox & L. Hautier) Ch. 18, 478–509 (Cambridge University Press, 2015).

39 Peterková, R., Lesot, H. & Peterka, M. Phylogenetic memory of developing mammalian dentition. J. Exp. Zool. B Mol. Dev. Evol. 306, 234–250 (2006).

40 Sadier, A. et al. Modeling Edar expression reveals the hidden dynamics of tooth signaling center patterning. PLoS Biol. 17, e3000064 (2019).

41 Hayden, L. et al. Developmental variability channels mouse molar evolution. Elife 9, e50103 (2020).

42 Kavanagh, K. D., Evans, A. R. & Jernvall, J. Predicting evolutionary patterns of mammalian teeth from development. Nature 449, 427–432 (2007).

43 Janis, C. M. & Fortelius, M. On the means whereby mammals achieve increased functional durability of their dentitions, with special reference to limiting factors. Biol. Rev. Camb. Philos. Soc. 63, 197–230 (1988).

44 Saarinen, J. & Lister, A. M. Fluctuating climate and dietary innovation drove ratcheted evolution of proboscidean dental traits. Nat. Ecol. Evol. 7, 1490–1502 (2023).

45 Davit-Béal, T., Tucker, A. S. & Sire, J. Y. Loss of teeth and enamel in tetrapods: fossil record, genetic data and morphological adaptations. J. Anat. 214, 477–501 (2009).

46 Charles, C., Solé, F., Rodrigues, H. G. & Viriot, L. Under pressure? Dental adaptations to termitophagy and vermivory among mammals. Evolution 67, 1792–1804 (2013).

47 Churchill, M. & Clementz, M. T. Functional implications of variation in tooth spacing and crown size in Pinnipedimorpha (Mammalia: Carnivora). The Anatomical Record 298, 878–902 (2015).

48 Peredo, C. M., Peredo, J. S. & Pyenson, N. D. Convergence on dental simplification in the evolution of whales. Paleobiology 44, 434–443 (2018).

49 Lafuma, F., Corfe, I. J., Clavel, J. & Di-Poï, N. Multiple evolutionary origins and losses of tooth complexity in squamates. Nat. Commun. 12, 6001 (2021).

50 Holliday, J. A. & Steppan, S. J. Evolution of hypercarnivory: the effect of specialization on morphological and taxonomic diversity. Paleobiology 30, 108–128 (2004).

51 Van Valkenburgh, B., Wang, X. & Damuth, J. Cope’s rule, hypercarnivory, and extinction in North American canids. Science 306, 101–104 (2004).

52 Robinson, B. W. & Wilson, D. S. Optimal foraging, specialization, and a solution to Liem’s paradox. Am. Nat. 151, 223–235 (1998).

53 Zachos, J., Pagani, M., Sloan, L., Thomas, E. & Billups, K. Trends, rhythms, and aberrations in global climate 65 Ma to present. Science 292, 686–693 (2001).

54 Lunt, D. J., Foster, G. L., Haywood, A. M. & Stone, E. J. Late Pliocene Greenland glaciation controlled by a decline in atmospheric CO2 levels. Nature 454, 1102–1105 (2008).

55 Ruddiman, W. F. & Kutzbach, J. E. Forcing of late Cenozoic northern hemisphere climate by plateau uplift in southern Asia and the American West. J. Geophys. Res. (D Atmospheres) 94, 18409–18427 (1989).

56 Maslin, M. A., Li, X., Loutre, M.-F. & Berger, A. The contribution of orbital forcing to the progressive intensification of Northern Hemisphere glaciation. Quat. Sci. Rev. 17, 411–426 (1998).

57 Bartoli, G. et al. Final closure of Panama and the onset of northern hemisphere glaciation. Earth Planet. Sci. Lett. 237, 33–44 (2005).

58 Willeit, M., Ganopolski, A., Calov, R. & Brovkin, V. Mid-Pleistocene transition in glacial cycles explained by declining CO2 and regolith removal. Sci. Adv. 5, eaav7337 (2019).

59 Gromov, I. & Polyakov, I. Y. Fauna of the USSR: Mammals. Vol. III, No 8 Voles (Microtinae) (Brill, Leiden, 1992).

60 Lazzari, V. et al. Mosaic convergence of rodent dentitions. PLoS One 3, e3607 (2008).

61 Rekovets, L. & Kovalchuk, O. Phenomenon in the evolution of voles (Mammalia, Rodentia, Arvicolidae). Vestn. Zool. 51, 99 (2017).

62 Maul, L. C., Masini, F., Parfitt, S. A., Rekovets, L. & Savorelli, A. Evolutionary trends in arvicolids and the endemic murid Mikrotia – New data and a critical overview. Quat. Sci. Rev. 96, 240–258 (2014).

63 Salazar-Ciudad, I. & Jernvall, J. How different types of pattern formation mechanisms affect the evolution of form and development. Evol. Dev. 6, 6–16 (2004).

64 Uller, T., Moczek, A. P., Watson, R. A., Brakefield, P. M. & Laland, K. N. Developmental bias and evolution: a regulatory network perspective. Genetics 209, 949–966 (2018).

65 Wagner, G. P. The influence of variation and of developmental constraints on the rate of multivariate phenotypic evolution. J. Evol. Biol. 1, 45–66 (1988).

66 Salazar-Ciudad, I. & Marín-Riera, M. Adaptive dynamics under development-based genotype–phenotype maps. Nature 497, 361–364 (2013).

67 Fortuna, M. A., Zaman, L., Ofria, C. & Wagner, A. The genotype-phenotype map of an evolving digital organism. PLoS Comp. Biol. 13, e1005414 (2017).

68 Greenbury, S. F., Louis, A. A. & Ahnert, S. E. The structure of genotype-phenotype maps makes fitness landscapes navigable. Nat. Ecol. Evol. 6, 1742–1752 (2022).

69 Escudé, É., Renvoisé, É., Lhomme, V. & Montuire, S. Why all vole molars (Arvicolinae, Rodentia) are informative to be considered as proxy for Quaternary paleoenvironmental reconstructions. J. Archaeol. Sci. 40, 11–23 (2013).

70 Lambert, J. E. & Rothman, J. M. Fallback foods, optimal diets, and nutritional targets: primate responses to varying food availability and quality. Annu. Rev. Anthropol. 44, 493–512 (2015).

71 Kropacheva, Y. E. & Zykov, S. V. An evaluation of individual seasonal changes in dental macro-and mesowear of wild-caught common vole (*Microtus arvalis* sensu lato) by the intravital impressions method. Mamm. Biol. 102, 501–516 (2022).

72 Machado, F. A. et al. Rules of teeth development align microevolution with macroevolution in extant and extinct primates. *Nat*. Ecol. Evol. 7, 1729–1739 (2023).

73 Felice, R. N., Randau, M. & Goswami, A. A fly in a tube: macroevolutionary expectations for integrated phenotypes. Evolution 72, 2580–2594 (2018).

74 Salazar-Ciudad, I. & Jernvall, J. Graduality and innovation in the evolution of complex phenotypes: insights from development. J. Exp. Zool. B Mol. Dev. Evol. 304, 619–631 (2005).

75 Benson, R. B. & Choiniere, J. N. Rates of dinosaur limb evolution provide evidence for exceptional radiation in Mesozoic birds. Proc. R. Soc. Lond., Ser. B: Biol. Sci. 280, 20131780 (2013).

76 Liu, S. Y. et al. A new vole from Xizang, China and the molecular phylogeny of the genus *Neodon* (Cricetidae: Arvicolinae). Zootaxa 3235, 1–22-21–22 (2012).

77 Upham, N. S., Esselstyn, J. A. & Jetz, W. Inferring the mammal tree: species-level sets of phylogenies for questions in ecology, evolution, and conservation. PLoS Biol. 17, e3000494 (2019).

78 Withnell, C. B. A new perspective on the systematics and paleobiogeography of arvicoline rodents and the first radiometric age of the fauna from Cumberland Bone Cave, Maryland Ph.D. thesis, The University of Texas at Austin (2020).

79 Agusti, J. The *Allophaiomys* complex in southern Europe. Geobios 25, 133–144 (1992).

80 Chaline, J., Laurin, B., Brunet-Lecomte, P. & Viriot, L. Morphological trends and rates of evolution in arvicolids (Arvicolidae, Rodentia): towards a punctuated equilibria/disequilibria model. Quat. Int. 19, 27–39 (1993).

81 Martin, R. D. in Evolution of Tertiary Mammals of North America Vol. 2 (eds C. M. Janis, G. F. Gunnell, & M. D. Uhen) 480–498 (Cambridge University Press, 2008).

82 Gibbard, P. L. & Head, M. J. The newly-ratified definition of the Quaternary System/Period and redefinition of the Pleistocene Series/Epoch, and comparison of proposals advanced prior to formal ratification. Episodes 33, 152–158 (2010).

83 Schneider, C. A., Rasband, W. S. & Eliceiri, K. W. NIH Image to ImageJ: 25 years of image analysis. Nat. Methods 9, 671–675 (2012).

84 Markova, E. Assessment of tooth complexity in arvicolines (Rodentia): a morphotype ranking approach. Biol. Bull. 41, 589–600 (2014).

85 Hinton, M. A. C. Monograph of the Voles and Lemmings (Microtinae) Living and Extinct. Vol. 1 (Trustees of the British Museum (Natural History), London, 1926).

86 Hibbard, C. W. Mammals of the Rexroad Formation from Fox Canyon, Kansas. Contr. Mus. Paleontol. Univ. Michigan 8, 113–192 (1950).

87 Jukes, T. H. & Cantor, C. R. in Mammalian Protein Metabolism (eds. H. N. Munro) Vol. 3, 21–132 (Academic Press, Cambridge MA, 1969).

88 Pagel, M. Detecting correlated evolution on phylogenies: a general method for the comparative analysis of discrete characters. Proc. R. Soc. Lond., Ser. B: Biol. Sci. 255, 37–45 (1994).

89 Lewis, P. O. A likelihood approach to estimating phylogeny from discrete morphological character data. Syst. Biol. 50, 913–925 (2001).

90 Revell, L. J. phytools: an R package for phylogenetic comparative biology (and other things). Methods Ecol. Evol. 3, 217–223 (2012).

91 Hurvich, C. M. & Tsai, C.-L. Regression and time series model selection in small samples. Biometrika 76, 297–307 (1989).

92 Venditti, C., Meade, A. & Pagel, M. Multiple routes to mammalian diversity. Nature 479, 393–396 (2011).

93 Baker, J., Meade, A., Pagel, M. & Venditti, C. Positive phenotypic selection inferred from phylogenies. Biol. J. Linn. Soc. 118, 95–115 (2016).

94 Plummer, M., Best, N., Cowles, K. & Vines, K. CODA: convergence diagnosis and output analysis for MCMC. R News 6, 7–11 (2006).

95 R Core Team. R: A language and environment for statistical computing. http://www.R-project.org/ (R Foundation for Statistical Computing, Vienna, Austria, 2024).

96 Xie, W., Lewis, P. O., Fan, Y., Kuo, L. & Chen, M.-H. Improving marginal likelihood estimation for Bayesian phylogenetic model selection. Syst. Biol. 60, 150–160 (2011).

97 Gelfand, A., Gilks, W., Richardson, S. & Spiegelhalter, D. Markov Chain Monte Carlo in Practice. (Chapman & Hall/CRC, 1996).

98 Yu, G., Smith, D. K., Zhu, H., Guan, Y. & Lam, T. T. Y. ggtree: an R package for visualization and annotation of phylogenetic trees with their covariates and other associated data. Methods Ecol. Evol. 8, 28–36 (2017).

99 Garnier, S. et al. viridis – Colorblind-friendly color maps for R. 10.5281/zenodo.4679424 (R package version 0.6.4, 2024).

100 Shmueli, G., Minka, T. P., Kadane, J. B., Borle, S. & Boatwright, P. A useful distribution for fitting discrete data: revival of the Conway–Maxwell–Poisson distribution. J. Roy. Stat. Soc. Ser. C. (Appl. Stat.) 54, 127–142 (2005).

101 Sellers, K. F. & Shmueli, G. A flexible regression model for count data. Ann. Appl. Stat., 943–961 (2010).

102 Calcagno, V. & de Mazancourt, C. glmulti: an R package for easy automated model selection with (generalized) linear models. J. Stat. Softw. 34, 1–29 (2010).

103 Brooks, M. E. et al. glmmTMB balances speed and flexibility among packages for zero-inflated generalized linear mixed modeling. R J. 9, 378–400 (2017).

104 Li, D., Dinnage, R., Nell, L. A., Helmus, M. R. & Ives, A. R. phyr: An r package for phylogenetic species-distribution modelling in ecological communities. Methods Ecol. Evol. 11, 1455–1463 (2020).

105 Pearson, K. X. On the criterion that a given system of deviations from the probable in the case of a correlated system of variables is such that it can be reasonably supposed to have arisen from random sampling. London Edinburgh Philos. Mag. & J. Sci. 50, 157–175 (1900).

106 Chambers, J. M. & Hastie, T. J. Statistical Models in S. (Wadsworth & Brooks/Cole Advanced Books & Software, 1992).

107 Dunn, P. K. & Smyth, G. K. Randomized quantile residuals. J. Comput. Graph. Stat. 5, 236–244 (1996).

108 Hartig, F. DHARMa: Residual Diagnostics for Hierarchical (Multi-Level / Mixed) Regression Models. https://CRAN.R-project.org/package=DHARMa (R package version 0.4.6, 2024).

109 Paradis, E. & Claude, J. Analysis of comparative data using generalized estimating equations. J. Theor. Biol. 218 175–185 (2002).

110 Ho, L. S. T. & Ané, C. A linear-time algorithm for Gaussian and non-Gaussian trait evolution models. Syst. Biol. 63, 397–408 (2014).

111 Ives, A. R. & Garland, T., Jr. Phylogenetic logistic regression for binary dependent variables. Syst. Biol. 59, 9–26 (2010).

112 Hosmer Jr, D. W., Lemeshow, S. & Sturdivant, R. X. Applied Logistic Regression. Vol. 398 (John Wiley & Sons, Hoboken NJ, 2013).

113 Lele, S. R., Keim, J. L. & Solymos, P. ResourceSelection: Resource Selection (Probability) Functions for Use-Availability Data. https://github.com/psolymos/ResourceSelection (R package version 0.3-6, 2024)

114 Metscher, B. D. MicroCT for comparative morphology: simple staining methods allow high-contrast 3D imaging of diverse non-mineralized animal tissues. BMC Physiol. 9, 11 (2009).

115 Stalling, D., Westerhoff, M. & Hege, H. in The Visualization Handbook (eds C. D. Hansen & C. R. Johnson) Ch. 38, 749–767 (Butterworth-Heinemann, Oxford, 2005).

116 Sahlberg, C., Mustonen, T. & Thesleff, I. Explant cultures of embryonic epithelium: analysis of mesenchymal signals. Methods Mol. Biol., 373–382 (2009).

