## Supplementary Information guide for "Six million years of vole dental evolution driven by tooth development"

This study is supported by the following supplementary information:

**Supplementary Data 1:** species-level dataset in CSV format including cusp numbers for the first lower molars (m1) of 151 extant and extinct Arvicolinae and one extinct cricetid outgroup and linear measurements of the m1 (length, width, length/width ratio) for 126 of these arvicoline species and their outgroup, with corresponding references. First and last appearance data cross-referenced based on sources cited in the Paleobiology Database (PBDB) are included when available.

**Supplementary Data 2:** time-calibrated phylogenetic tree of 151 species of Arvicolinae and their outgroup, in Newick format.

**Supplementary Data 3:** time-calibrated phylogenetic tree of 109 species of Arvicolinae bearing a seven- or nine-cusped m1 of known dimensions, in Newick format.

**Supplementary Data 4:** time-calibrated phylogenetic tree of 126 species of Arvicolinae with known m1 cusp number and dimensions and their outgroup, in Newick format.

**Supplementary Data 5:** dataset in CSV format of cusp numbers and linear measurements (length, width, length/width ratio) for explants of bank vole m1 germs developing *in vitro*.
